# Deciphering subcellular localization-dependent functions of Hog1 MAPK in *Cryptococcus neoformans*

**DOI:** 10.64898/2025.12.17.694976

**Authors:** Yu-Byeong Jang, Yong-Sun Bahn

## Abstract

The Hog1 mitogen-activated protein kinase regulates stress adaptation, sexual differentiation, and virulence traits by dynamically shuttling between the cytoplasm and nucleus in the fungal pathogen *Cryptococcus neoformans*, a leading cause of fatal fungal meningoencephalitis worldwide. However, how spatial partitioning governs Hog1 pathway specificity remains poorly understood. Here, we generated genetically engineered strains expressing localization-restricted Hog1 variants including membrane-tethered and constitutively nuclear, and we compared their phenotypic traits with those of a fully functional Hog1–mCherry strain. The membrane-tethered Hog1 restored thermotolerance and antifungal resistance in the *hog1*Δ mutant, whereas nuclear Hog1 was necessary for osmotic and endoplasmic reticulum (ER) stress response and for capsule and melanin biosynthesis. These localization-specific effects were corroborated by the expression profiles of genes involved in glycerol biosynthesis, under osmotic stress or fludioxonil treatment. Measurement of intracellular glycerol revealed that the plasma membrane-tethering of Hog1 caused excessive accumulation, underscoring the importance of spatiotemporal regulation of Hog1 activity. Also, nuclear Hog1 uniquely reinstated osmoadaptation and conferred tunicamycin tolerance independent of canonical Ire1-Hxl1 splicing, suggesting a nucleus-centric module for ER stress protection. Conversely, membrane-tethered Hog1 enhanced glycerol accumulation and restored amphotericin B resistance while increasing azole susceptibility despite normal ergosterol levels, implicating localization-biased regulation upstream of sterol abundance. In the developmental program, membrane-tethered Hog1 dominantly suppressed the Cpk1-mediated mating. Collectively, these findings demonstrate that Hog1 compartmentalization is a key determinant of stress adaptation, antifungal resistance, differentiation, and virulence regulation in *C. neoformans*.

**Summary:** This study elucidates the spatial regulation of Hog1 MAPK in *Cryptococcus neoformans* using membrane-tethered (CAAX) and constitutively nuclear (NLS) mutants. Unlike model yeasts, *C. neoformans* requires precise Hog1 compartmentalization for distinct functions: nuclear translocation is essential for osmotic and ER stress adaptation, while cytoplasmic retention is critical for regulating glycerol homeostasis and repressing sexual differentiation. Notably, membrane-tethered Hog1 causes toxic glycerol accumulation and azole hypersensitivity, suggesting a metabolic maintenance where nuclear translocation prevents cytoplasmic hyperactivation. These findings define a specific spatial function for Hog1, highlighting evolutionary divergence in fungal stress signaling and virulence regulation.

## Introduction

The high-osmolarity glycerol (HOG) pathway is an evolutionarily conserved signaling cascade in eukaryotes that plays a pivotal role in sensing and responding to diverse environmental stresses, particularly osmotic stress (de Nadal and Posas 2022). First characterized in the model yeast *Saccharomyces cerevisiae*, the HOG pathway is essential for osmoadaptation in this organism (Brewster et al. 1993; Canovas and Nebreda 2021). Because stress sensing and adaptation are critical for the survival of fungal pathogens within hostile host environments, the Hog1 MAPK pathway has been extensively investigated in fungi infecting both plants and animals (Bahn 2008; Day and Quinn 2019; Jiang et al. 2018). Beyond its role in stress adaptation, the HOG pathway also contributes to fungal differentiation, antifungal drug resistance, and virulence.

The primary signaling pathway upstream of the fungal Hog1 MAPK module is a multicomponent phosphorelay system comprising hybrid histidine kinases (HHKs), a histidine-containing phosphotransfer protein (HPt), and response regulators (RRs). In *S. cerevisiae*, under isotonic conditions, the sole HHK, Sln1, undergoes continuous autophosphorylation and sequentially transfers its phosphate group to the Hpt protein Ypd1 and then to the response regulator Ssk1. Phosphorylated Ssk1 remains inactive and does not trigger downstream signaling. Upon exposure to hyperosmotic stress, Sln1 becomes dephosphorylated, halting the phosphorelay to Ypd1 and Ssk1. Consequently, unphosphorylated Ssk1 activates the MAPK kinase kinase (MAP3K) Ssk2/22, initiating a phosphorylation cascade through the MAPK kinase (MAP2K) Pbs2 and ultimately activating Hog1 MAPK. Activated Hog1 then translocates to the nucleus, where it induces the transcription of osmoprotective genes, or directly regulates metabolic enzymes that elevate intracellular glycerol levels, thereby facilitating adaptation to osmotic stress (de Nadal and Posas 2022).

The core signaling architecture of the HOG pathway is broadly conserved among fungal species, with the major point of divergence being the number of hybrid histidine kinases (HHKs). Unlike *S. cerevisiae*, which possesses a single HHK (Sln1), most fungi contain multiple HHKs, including 11 in *Neurospora crassa*, 13 in *Aspergillus fumigatus*, 15 in *Aspergillus nidulans*, and 21 in *Zymoseptoria tritici* (Banno et al. 2007; Hayashi et al. 2014; Herivaux et al. 2016). The specific contributions of these HHKs to HOG pathway regulation remain incompletely defined. Nevertheless, as most fungi harbor only one HPt protein, it is plausible that signals from multiple HHKs converge at this common node. Such convergence may allow fine-tuned modulation of the HOG pathway in response to diverse environmental stimuli.

*Cryptococcus neoformans*, a basidiomycetous fungal pathogen that causes fatal meningoencephalitis in immunocompromised individuals, exemplifies a species harboring multiple HHKs, designated Tco1 through Tco7. Among these, Tco1, which contains a HAMP domain, and Tco2, which possesses dual HHK domains, are implicated in HOG pathway signaling. This is supported by the observation that deletion of both *TCO1* and *TCO2* confers complete resistance to the fungicide fludioxonil, phenocopying the *HOG1* deletion mutant. Downstream of Tco1/2, a single HPt (Ypd1) and two RRs (Ssk1 and Skn7) are present (Bahn et al. 2006). These components transmit signals through the MAPK cascade composed of Ssk2 (MAP3K), Pbs2 (MAP2K), and Hog1 (MAPK) (Bahn and Jung 2013). In *C. neoformans*, the HOG pathway regulates not only stress response and antifungal resistance but also sexual differentiation, capsule and melanin production, and virulence (Bahn et al. 2005).

A distinctive feature of the cryptococcal HOG pathway is the variation in basal Hog1 phosphorylation among pathogenic *Cryptococcus* species (Bahn et al. 2005). In the H99 strain (*C. neoformans* reference genome strain), Hog1 is constitutively phosphorylated under basal conditions and undergoes dephosphorylation upon exposure to environmental stress, such as osmotic shock (Bahn et al. 2005). In contrast, in the JEC21 strain (*C. deneoformans* reference genome strain), Hog1 remains largely unphosphorylated under unstressed conditions but is rapidly phosphorylated in response to stress, resembling the pattern observed in *S. cerevisiae* (Bahn et al. 2005). These opposing phosphorylation dynamics are associated with distinct subcellular localization patterns: in H99, Hog1 is enriched in the nucleus even in the absence of stress. This elevated nuclear accumulation may underlie the enhanced stress resistance of H99 compared with JEC21 (Bahn et al. 2005).

Despite extensive research on the fungal HOG signaling pathway, the regulatory mechanism underlying Hog1 MAPK function remains incompletely understood. In *S. cerevisiae*, Hog1 resides in the cytoplasm in its dephosphorylated state under basal conditions and translocates to the nucleus upon osmotic stress, where it activates transcription of downstream effector genes (Ferrigno et al. 1998; Huang et al. 2020). Intriguingly, this nuclear translocation appears to be dispensable for Hog1-mediated stress resistance, as membrane-tethered Hog1 can fully restore stress tolerance in the *hog1*Δ mutant (Westfall et al. 2008). A similar observation has been reported in *Candida albicans*, where membrane-tethered Hog1 likewise rescues the *hog1*Δ mutant phenotype (Day et al. 2017). Nevertheless, important differences exist between the two ascomycete yeasts. In *S. cerevisiae*, membrane-tethered Hog1 fails to induce expression of target genes, whereas in *C. albicans*, it retains transcriptional activity (Day et al. 2017; Westfall et al. 2008). Moreover, in *C. albicans*, the *hog1*Δ mutant can be fully complemented by either a membrane-tethered or constitutively nuclear Hog1, indicating that Hog1 remains functional regardless of its subcellular location (Day et al. 2017).

In this study, we investigated whether Hog1 functionality in the basidiomycetous yeast *C. neoformans* depends on its subcellular localization. Specifically, we asked whether Hog1 requires nuclear translocation for its activity or, as in ascomycete yeasts, can function when restricted to the plasma membrane. To address this question, we generated two engineered *C. neoformans* strains: one expressing Hog1 fused at the C-terminus with the Ras1-CAAX motif for plasma membrane anchoring, and another expressing Hog1 tagged at the N-terminus with a nuclear localization signal (NLS) for constitutive nuclear targeting. Our findings reveal that, in *C. neoformans*, Hog1 exerts distinct and complementary functions that depend on its subcellular localization.

## Materials and Methods

### Construction of plasma membrane-tethered and nuclear Hog1 mutants

The *C. neoformans hog1*Δ mutant (YSB64) (Bahn et al. 2005) was used as a parental strain to newly generate three complemented strains in this study: *hog1*Δ::*HOG1-mCherry* (Hog1-mCh), *hog1*Δ::*HOG1-mCherry-CAAX* (Hog1-CAAX), and NLS-Hog1 (*hog1*Δ::*NLS*_SV40_*-HOG1-mCherry*) (Table S1). For strain construction, the native *HOG1* promoter and entire ORF was subcloned into the pNEO_mCherry vector as previously described (So et al. 2017). The *HOG1-mCherry-CAAX* and *NLS_SV40_-HOG1-mCherry* plasmids were engineered to generate plasma membrane tethered- and constitutively nuclear Hog1 variants, respectively. Linearized plasmids were introduced into the *hog1*Δ mutant by biolistic transformation. The parental strain was cultured overnight in 50 mL YPD (yeast extract-peptone-dextrose) medium at 30°C with shaking at 200 rpm. Cells were harvested, resuspended in 5 mL fresh YPD, spread onto YPD agar containing 1 M sorbitol, and incubated for 3 h at 30°C. Plasmid DNA was coated onto 0.6 µm gold bead (600 µg; Bio-Rad Laboratories) using 2.5 M CaCl_2_ and 2 µL spermidine (S-0266, Sigma Aldrich) and delivered into cells using a particle delivery system (PDS-1000, Bio-Rad Laboratories) at 1500 psi. After 4 h of recovery at 30°C, cells were scraped and plated on YPD agar containing G418 (40 µg/mL) for selection. Transformants were verified by diagnostic PCR, and the size of Hog1 fusion proteins were confirmed by western blot analysis.

### Protein extraction and immunoblotting assay

The wild-type, Hog1-mCh, Hog1-CAAX, and NLS-Hog1 strains were cultured overnight in YPD medium at 30°C. Cells were diluted to an optical density at 600 nm (OD_600_) of 0.2 in fresh YPD and incubated to exponential phase (OD_600_ ≍ 0.8) at 30°C. Early exponential phase samples were centrifuged and immediately frozen in liquid nitrogen. Frozen cells were resuspended in 0.7 mL of lysis buffer containing 50 mM Tris pH 7.5, 1% sodium deoxycholate, 5 mM sodium pyrophosphate, 0.1 µM sodium orthovanadate, 50 mM sodium fluoride, 0.1% SDS, 1% Triton X-100, 0.5 µM PMSF, and 1x protease inhibitor cocktail (Set IV; Calbiochem). Cells were disrupted using a bead beater (FastPrep-24™ 5G; MP Biomedicals) at 6.0 m/s for 30 sec, with 10 cycles applied to enhance extraction efficiency, particularly for membrane-associated Hog1 variants. Samples were then centrifuged at 13,000 rpm for 20 min at 4°C. The supernatants were then transferred to new tubes. Protein concentrations were determined using the Pierce™ BCA Protein Assay Kits (ThermoFisher Scientific) with bovine serum albumin (BSA) as the standard. Equalized protein samples were mixed with 5x sample buffer (CBS002; LPS solution), heated to 95°C for 15 min, and stored at -20°C until use. Immunoblotting was performed using the following primary antibodies: phospho-p38 MAPK antibody (#9212, 2000:1; Cell signaling), anti-Hog1 (2000:1; custom antibody), and mCherry polyclonal antibody (PA5-34974, 1000:1; Invitrogen). The secondary antibody was HRP-conjugated mouse anti-rabbit IgG (sc-2357, 4000:1; Santa Cruz Biotechnology). HRP signals were visualized using an enhanced chemiluminescence (ECL) detection system (D-Plus ECL Pico Alpha System), and images were captured with the ChemiDoc XRS^+^ (Bio-Rad Laboratories).

### Visualization of mCherry-fused protein

The mCherry-tagged Hog1 strains were cultured overnight in YPD medium at 30°C. Cultures were diluted to an OD_600_ of 0.2 in fresh YPD and incubated to mid-log phase (OD_600_ ≍ 0.8) at 30°C. To induce osmotic stress, NaCl was added to a final concentration of 1 M, and samples were collected at 0, 10, and 60 min post-treatment. Cells were harvested, washed with phosphate-buffered saline (PBS), and stained with Hoechst dye for 30 min in the dark at room temperature. Excess dye was removed by two additional PBS washes. The stained cells were mounted on glass slides and imaged using a Zeiss LSM900 confocal laser scanning microscope mounted on an Axio Observer Z1 inverted microscope (Carl Zeiss) equipped with a Plan-Apochromat 100×/1.4 NA oil-immersion objective lens. mCherry fluorescence was excited with a 561 nm laser line and detected at 565–700 nm, Hoechst-stained nuclei were excited with a 405 nm laser line and detected at 400– 495 nm, and cell morphology was visualized using the transmitted-light photomultiplier tube (T-PMT) detector. Image acquisition and quantitative line-scan fluorescence intensity analysis (used to calculate Pearson’s correlation coefficients between mCherry and Hoechst signals presented in Fig. 1d) were performed using ZEN 3.0 imaging software (Carl Zeiss).

**Fig. 1.**
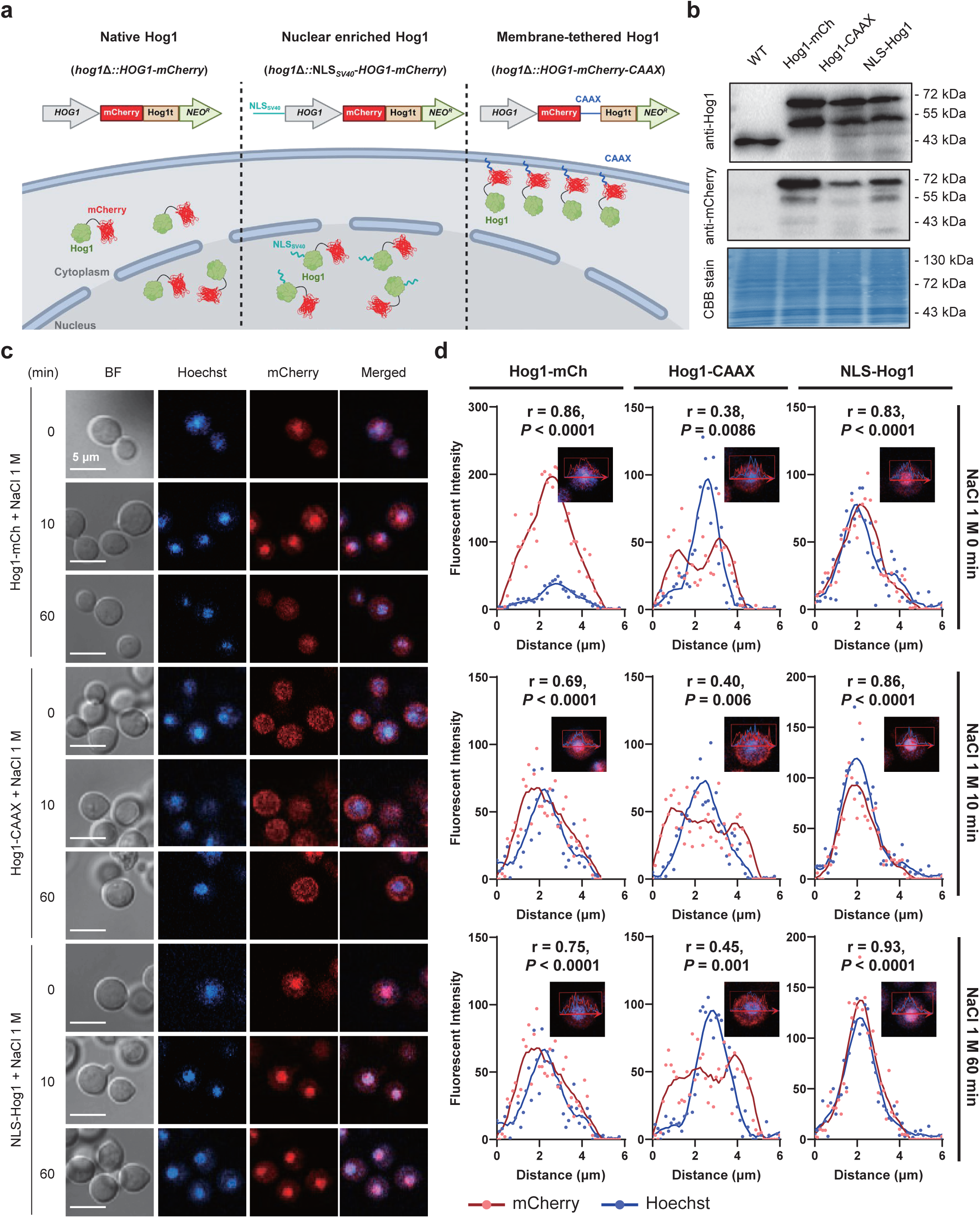
Construction of membrane-tethered and nuclear-enriched Hog1 mutant and confirmation of their subcellular localization. (a) The construction of Hog1-mCh, Hog1-CAAX, and NLS-Hog1 mutants was illustrated using BioRender (https://BioRender.com/v91v944). (b) The protein expression of wild-type and all three Hog1-tagged mutants confirmed by western blot analysis using anti-Hog1 and anti-mCherry antibodies. Coomassie Brilliant Blue (CBB) staining of the membrane was used as a loading control. The predicted molecular weights of each protein are as follows: native Hog1, 41.2 kDa; Hog1-mCh, 68 kDa; Hog1-CAAX, 75 kDa; and NLS-Hog1, 78.3 kDa. The ∼43 kDa band detected in the WT lane corresponds to endogenous native Hog1, which is absent in the mCherry-tagged Hog1 strains due to deletion of the endogenous *HOG1* gene. The ∼55 kDa band detected in the Hog1-mCh, Hog1-CAAX, and NLS-Hog1 samples by both anti-Hog1 and anti-mCherry antibodies is presumed to represent a proteolytic cleavage product of the Hog1-mCherry fusion proteins, as this band is absent in the wild-type strain and retains portions of both the Hog1 and mCherry moieties. (c) Subcellular localization of Hog1 in each of the three mutants. *C. neoformans* cells expressing Hog1-mCh, Hog1-CAAX, or NLS-Hog1 were synchronized to mid-exponential phase and treated with 1 M NaCl for 0 (basal, pre-stress), 10, or 60 min, then stained with Hoechst 33342 to visualize nuclei. Cells were imaged using a Zeiss LSM900 confocal laser scanning microscope as described in Materials and Methods. Rows correspond to the three Hog1 variants under sequential osmotic stress time points (top to bottom: Hog1-mCh + NaCl 1 M, Hog1-CAAX + NaCl 1 M, NLS-Hog1 + NaCl 1 M, each at 0, 10, and 60 min). Columns show (left to right): brightfield (BF), Hoechst (blue, nuclear), mCherry (red, Hog1 variants), and merged channels (blue and red). Scale bar, 5 µm. (d) Quantitative line-scan analysis of mCherry and Hoechst fluorescence intensities along the indicated lines in the corresponding cells in Fig. 1c. Pearson’s correlation coefficients (r) between the mCherry and Hoechst signals are shown for each cell, with *P* values from two-tailed correlation tests. High r values indicate strong spatial overlap between the two signals (i.e., nuclear enrichment of mCherry), whereas low r values indicate spatial separation (i.e., nuclear exclusion). The raw fluorescence intensity data underlying these line-scan profiles are provided in Supplementary Table S3.

### Quantitative reverse transcription-PCR (qRT-PCR) analysis

Wild-type, *hog1*Δ, Hog1-mCh, Hog1-CAAX, and NLS-Hog1 strains were inoculated into 50 mL of YPD broth and cultured overnight at 30°C. The cultures were then diluted into fresh YPD medium and grown to an OD_600_ ≍ 0.8, followed by treatment with 1 M NaCl, 10 µg/mL fludioxonil, or 3 mM hydrogen peroxide for the indicated times. The cells were harvested by centrifugation and lyophilized. The dried cells were disrupted using 3 mm glass bead, and total RNA was extracted using an RNA extraction kit (iNtRON Biotechnology). RNA concentration was quantified using a NanoDrop spectrophotometer (Bio-Rad Laboratories, USA). Complementary DNA (cDNA) was synthesized from 5 µg of total RNA. Quantitative reverse transcription-PCR (qRT-PCR) was performed on a CFX96 Real-Time PCR Detection System (Bio-Rad Laboratories) using gene-specific primers and TB Green (ThermoFisher Scientific). Relative transcript levels were calculated with the 2^−ΔΔCT^ method and normalized to *ACT1* expression. All data were illustrated using GraphPad Prism 10.0.

### Quantification of cell swelling and intracellular glycerol content

Wild-type, *hog1*Δ, Hog1-mCh, Hog1-CAAX, and NLS-Hog1 strains were cultured overnight in YPD medium. The cultures were diluted to an OD_600_ of 0.2 and further incubated for 4–5 h at 30°C until reaching an OD_600_ ≍ 0.8. Fludioxonil was then added to a final concentration of 2 or 10 μg/mL, and the cultures were incubated at 30°C for one or two days until reaching the stationary phase. After two days, the cells were harvested, washed with PBS, and observed under an optical microscope with 1000x magnification. The diameters of 50 randomly selected cells were measured to quantify cell swelling, and the data were visualized using GraphPad 10.0. For intracellular glycerol measurement, the same strains were cultured overnight, diluted to OD_600_ = 0.2, and further incubated 24 h. Then cells were then harvested, washed once with PBS, and boiled at 100°C for 10 min. The lysates were centrifuged at 13,000 rpm for 10 min, and the supernatants were used to determine intracellular glycerol levels using a Glycerol Assay Kit (MAK117, Sigma Aldrich).

#### Chemical and physical stress susceptibility assays

To evaluate osmotic stress tolerance, antifungal drug susceptibility, cell membrane integrity, oxidative stress responses, and thermosensitivity of the Hog1 variants, spotting assays were performed. Wild-type (H99), *hog1*Δ, Hog1-mCh, Hog1-CAAX, and NLS-Hog1 strains were cultured overnight in YPD medium at 30°C. The cultures were serially diluted 10-fold (10^-4^ to undiluted cells) and spotted on YPD agar plates. The plates were incubated under various conditions: different temperatures (30°C, 39°C, and 40°C), or YPD supplemented with osmotic stressors (NaCl, KCl, sorbitol), antifungal agents (amphotericin B, fludioxonil, fluconazole, voriconazole, ketoconazole), ER stress inducer (tunicamycin), oxidative stressors (H_2_O_2_, *tert*-butyl-BOOH, diamide, menadione), or membrane stress agent (SDS). For freeze-thaw stress test, cells were frozen in liquid nitrogen and thawed in 37°C water bath for 1 min. After incubation for 2-5 days, colony growth was imaged using GelDoc XR+ gel documentation system (Bio-Rad Laboratories).

#### *HXL1* splicing assay

Wild-type, *hog1*Δ, Hog1-mCh, Hog1-CAAX, and NLS-Hog1 strains were cultured overnight and diluted to an OD_600_ = 0.2. The cultures were then incubated for two days to reach the stationary phase, followed by treatment with 4 μg/mL tunicamycin. Cells were harvested at 0, 1, and 2 h post-treatment by centrifugation at 3,000 rpm for 10 min and subsequently lyophilized. Total RNA was extracted from the freeze-dried cells, and cDNA was synthesized as described above. To assess *HXL1* (CNAG_06134, Hac1/Xbp1-like gene 1) mRNA splicing, RT-PCR was performed using gene-specific primers for *HXL1* and *ACT1* (Appendix 2), PCR products were separated on 2% agarose gels, and band patterns were visualized and quantified using the GelDoc Imaging System (Bio-Rad Laboratories).

#### Ergosterol quantification

Ergosterol content was determined following previously described protocols with minor modifications. Wild-type and Hog1 variant strains were cultured in 100 mL of YPD medium at 30°C for 24 h. Each culture was then divided into two 50 mL aliquots, transferred to conical tubes, and lyophilized. One aliquot was weighed to determine dry cell mass for normalization, while the other was mixed with 5 mL of 25% alcoholic KOH solution and vortexed. The mixture was incubated at 80°C for 1 h, after which 3 mL of heptane and 1 mL of distilled water were added and vortexed to extract sterols. The heptane layer (200 μL) was diluted fivefold with 100% ethanol (800 μL), and ODs were measured at 281.5 nm and 230 nm. The percentage of ergosterol was calculated by subtracting the contribution of 24(28)-dehydroergosterol from the total absorbance at 281.5 nm, normalized to the dry pellet weight. The ergosterol content was determined using the following equation: % ergosterol = [(OD_281.5_ /290) × *F* / pellet weight] − [(OD_230_/518) × *F* / pellet weight], where *F* represents the ethanol dilution factor, and 290 and 518 are the E values (in % per centimeter) for crystalline ergosterol and 24(28)-dehydroergosterol, respectively.

#### Mating assay

To assess mating efficiency, the *MAT*a wild-type strain (H99) and its Hog1 mutants (*hog1*Δ, Hog1-mCh, Hog1-CAAX, and NLS-Hog1) were used together with the *MAT***a** wild-type strain (YL99**a**). All strains were cultured overnight in YPD medium at 30°C and washed twice with PBS. For mating analysis, *MAT*a wild-type or Hog1 mutant cells were mixed with *MAT***a** wild-type cells at a density of 10^8^ cells/mL each. The mixtures were spotted onto V8 mating medium (pH 7.0) or MS medium (pH 5.8) and incubated in the dark at 25°C for 1–2 weeks. Filamentation and sporulation were examined using DIC microscope (ECLIPSE Ni, Nikon) equipped with a DS-Qi2 camera (Nikon), or an Olympus BX51 microscope equipped with a SPOT Insight digital camera (Diagnostic Instruments, Inc).

#### Virulence factor production assays

Melanin induction assays were performed as previously described (Lee et al. 2019). Briefly, wild-type and Hog1 variant strains were cultured overnight at 30°C, harvested, washed three times with PBS, and resuspended in PBS. Each strain (3 µL) was spotted on Niger seed agar, epinephrine agar, and dopamine agar medium containing limited glucose (0.2%). The plates were incubated at 37°C and photographed for 1 to 3 days using a stereomicroscope (SMZ-168; Motic).

For capsule production, the same set of strains was grown overnight in YPD medium, harvested, and washed twice with PBS. Cell suspensions (3 µL) were spotted on Dulbecco’s Modified Eagle’s medium (DMEM) agar plates and incubated at 37°C for 2 days. Capsules were visualized by India ink staining (Remel Inc.), and fifty randomly selected cells were observed under a bright-field microscope. Capsule and cell body diameters were measured using Nikon NIS software to calculate capsule thickness (Jang et al. 2022).

Urease production was evaluated using a modified rapid urea hydrolysis (RUH) as described previously (Kwon-Chung et al. 1987). Wild-type and Hog1 variant strains were cultured overnight in YPD at 30°C, washed twice with distilled water, and adjusted to 2 × 108 cells/mL. Equal volumes (1 mL each) of cell suspension and 2x RUH broth (pH 6.8, containing urea, yeast extract, phenol red, and phosphate buffers) were mixed on ice to yield a final cell concentration of 1 × 108/mL. The mixtures were incubated at 37°C with shaking, and color changes were monitored hourly for up to 2 h. A magenta-red color indicated positive urease activity, whereas an orange-yellow color indicated a negative result. Borderline cases were quantified by measuring OD_560_, with values > 0.3 considered positive.

## Results

### Construction of plasma membrane-tethered and constitutively nuclear-localized Hog1 strains in *C. neoformans*

To investigate the functional significance of Hog1 subcellular localization in *C. neoformans*, we first generated a control complemented strain, *hog1*Δ::*HOG1-mCherry* (Hog1-mCh) (Fig. 1a; Supplementary Fig. S1a), in which a wild-type *HOG1* allele tagged at the C-terminus with mCherry was reintegrated at the native locus. Western blot analysis using anti-mCherry and anti-Hog1 antibodies confirmed the expression and predicted molecular weight (68 kDa) of the Hog1-mCherry fusion protein (Fig. 1b; Supplementary Fig. S1b; Supplementary Fig. S3a). This complemented strain exhibited wild-type phenotypes across all Hog1-dependent traits (Supplementary Fig. S1c, S1d), indicating that C-terminal mCherry tagging does not impair Hog1 function. Fluorescence microscopy revealed that native Hog1 localized to both the cytoplasm and nucleus, with a prominent nuclear enrichment under basal, pre-stress condition (time 0) (Fig. 1c), consistent with previous observations using a Hog1-GFP fusion (Bahn et al. 2005). Upon osmotic stress (1 M NaCl), Hog1 transiently accumulated more in the nucleus at early time points before equilibrating between the nucleus and cytoplasm (Fig. 1c), in agreement with prior findings (Bahn et al. 2005).

To construct a plasma membrane-tethered Hog1 strain, we engineered the *HOG1-mCherry-CAAX* allele by fusing the C-terminus of *HOG1*-*mCherry* in-frame with the C-terminal CAAX palmitoylation motif (CCRGCVVL) derived from cryptococcal Ras1 (Nichols et al. 2009). The CAAX motif—comprising a cysteine (C), two aliphatic amino acids (AA), and an amino acid (X)—undergoes post-translational modifications that anchor proteins to the plasma membrane (Wright and Philips 2006). In *C. neoformans*, Ras1 is known to localize to the plasma membrane through this dual CAAX palmitoylation motif (Nichols et al. 2009). Targeted integration of *HOG1-mCherry-CAAX* into the *hog1*Δ background yielded the *hog1*Δ::*HOG1-mCherry-CAAX* (Hog1-CAAX) strain (Fig. 1a). Western blot analysis confirmed the correct expression and predicted size (75 kDa) of the Hog1-CAAX fusion (Fig. 1b; Supplementary Fig. S3a). As expected, Hog1-CAAX was strongly enriched at the plasma membrane and remained membrane-associated following osmotic stress (Fig. 1c).

To generate a strain with constitutive nuclear localization, we fused a nuclear localization signal (NLS) from simian virus 40 (NLS_SV40_, PKKKRKV) to the N-terminus of *HOG1-mCherry*. The NLS_SV40_ sequence is widely used in yeast genetics to drive nuclear targeting (Bordonne 2000; Hahn et al. 2008) and has previously been employed in *C. albicans* to constitutively localize Hog1 to the nucleus (Day et al. 2017). The resulting *hog1*Δ::*NLS_SV40_*-*HOG1-mCherry* (NLS-Hog1) strain was generated by reintegrating the *NLS _SV40_*-*HOG1-mCherry* construct at the native Hog1 locus within the *hog1*Δ mutant (Fig. 1a). Western blot analysis confirmed protein expression and the expected molecular weight (78.3 kDa) using anti-Hog1 and anti-mCherry antibodies (Fig. 1b; Supplementary Fig. S3a). Fluorescence microscopy revealed strong nuclear enrichment of NLS-Hog1 under both basal and stress conditions, indicating stable nuclear localization (Fig. 1c).

To quantitatively validate the qualitative localization patterns observed in Fig. 1c, we performed line-scan analysis of mCherry and Hoechst fluorescence intensities along defined transects through representative cells (Fig. 1d). Consistent with the visual patterns, both the Hog1-mCh and NLS-Hog1 strains exhibited strong spatial correlation between mCherry and Hoechst signals, confirming predominant nuclear enrichment. Notably, NLS-Hog1 displayed consistently higher Pearson’s correlation coefficients (r = 0.83–0.93) than Hog1-mCh (r = 0.69–0.86) across all time points, with the correlation remaining stable following osmotic stress. This indicates that the SV40 NLS drives more robust and stress-independent nuclear retention, consistent with its intended role as a constitutive nuclear-targeting signal. In contrast, the Hog1-CAAX strain showed markedly reduced correlation (Pearson’s r = 0.38–0.45), consistent with its nuclear exclusion and plasma membrane tethering.

### Membrane-tethered and nuclear Hog1 independently regulate transcriptional activation of effector genes for osmoadaptation

Previous studies in *S. cerevisiae* and *C. albicans* have shown that both plasma membrane-tethered Hog1 and nuclear-enriched Hog1 variants can restore osmotic resistance in *hog1*Δ mutants (Alepuz et al. 2001; Day et al. 2017; Shiraishi et al. 2018; Westfall et al. 2008). These findings suggest that, in these ascomycete yeasts, Hog1 contributes to osmoadaptation through two distinct mechanisms: transcriptional activation of stress-responsive effector genes in collaboration with nuclear-localized transcription factors and direct cytoplasmic regulation of metabolic enzymes involved in glycerol biosynthesis. Alternatively, membrane-tethered Hog1 may activate cytoplasmic transcription factors, which can subsequently undergo nuclear translocation and induce effector genes for osmoadaptation.

To determine whether Hog1 exerts a similar dual role in *C. neoformans*, we examined whether localization-restricted Hog1 could restore osmotic resistance in the *hog1*Δ mutant. The nuclear Hog1 variant restored wild-type resistance levels, whereas the membrane-tethered Hog1 variant exhibited marked hypersensitivity to osmotic stress (Fig. 2a). Strikingly, the Hog1-CAAX strain was even more osmosensitive than the *hog1*Δ mutant under glucose-deprived conditions (Fig. 2a). These indicate that nuclear localization of Hog1 is indispensable for osmoadaptation in C. neoformans.

**Fig. 2.**
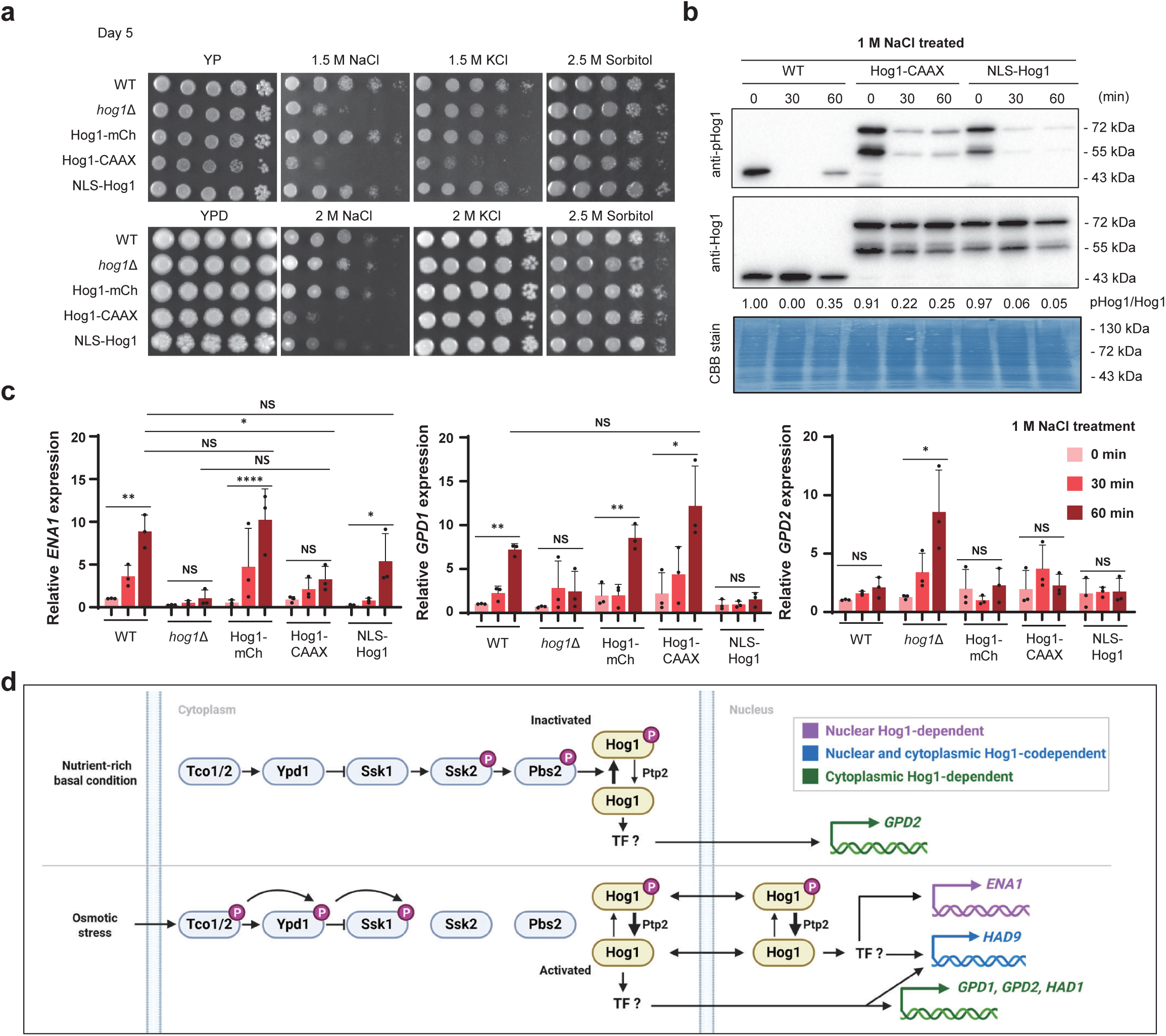
Cytoplasmic Hog1 mutant showed hypersensitive to osmotic stress. (a) Phenotypic assay of wild-type (WT), *hog1*Δ, Hog1-mCh, Hog1-CAAX, and NLS-Hog1 mutants on YP or YPD media supplemented with osmotic stresses (NaCl, KCl, or sorbitol). Ten-fold serial dilutions (10^-4^ to undiluted cells) were spotted and incubated for 5 days at 30°C. (b) Western blot analysis demonstrating the phosphorylation and total protein levels of Hog1 in WT, Hog1-CAAX, and NLS-Hog1 mutants upon exposure to 1 M NaCl. Hog1 phosphorylation was detected using a phospho-p38 antibody, and total Hog1 levels were detected using an anti-Hog1 antibody. Coomassie Brilliant Blue (CBB) staining of the membrane was used as a loading control. The ∼43 kDa band in the WT lane corresponds to endogenous native Hog1, which is absent in the Hog1-CAAX and NLS-Hog1 strains. The ∼55 kDa band in the Hog1-CAAX and NLS-Hog1 lanes represents a proteolytic cleavage product of the Hog1-mCherry fusion proteins (see Fig. 1b for characterization). (c) Real-time quantitative reverse transcription PCR (qRT-PCR) analysis of Hog1-regulated osmotic stress-responsive genes (*ENA1, GPD1,* and *GPD2*) after treatment with 1 M NaCl for the indicated times. The expression level of each gene is presented as relative fold-change normalized to the *ACT1* gene expression and untreated control. (d) Graphical summary of Hog1-dependent transcriptional regulation under osmotic stress illustrated by Biorender (https://BioRender.com/37dsts7). Transcriptional programs that depend on nuclear Hog1 (pink), on both nuclear and cytoplasmic Hog1 (blue), or on cytoplasmic Hog1 (green) are likely mediated by transcription factors under hyperosmotic conditions. Expression data for *ENA1*, *GPD1*, and *GPD2* are shown in panel (c); expression data for *HAD1*, *HAD6*, *HAD8*, and *HAD9* are presented in Supplementary Fig. S4a.

Despite these contrasting stress phenotypes, Hog1 phosphorylation dynamics were comparable among Hog1-CAAX, NLS-Hog1, and the wild-type strains. Both localization variants exhibited similar dephosphorylation kinetics following exposure to 1 M NaCl, although their basal phosphorylation levels were lower than those of the wild-type strain. Interestingly, we observed that both membrane-tethered and nuclear Hog1 undergo dephosphorylation upon osmotic stress, but Hog1-CAAX sustained the phosphorylation signal slightly longer than WT or NLS-Hog1 (Fig. 2b). This suggests the involvement of distinct phosphatases acting in the cytoplasmic and nuclear compartments. These subtle phosphorylation/dephosphorylation kinetic differences may partially account for their phenotypic differences.

Subsequently, we examined the expression of known osmotic stress-inducible effector genes in the Hog1 localization variants. In *C. neoformans*, Ena1 (encoding a Na^+^/ATPase) functions as a key cation transporter required for osmoadaptation and is strongly upregulated upon osmotic stress in a Hog1-dependent manner (Jung et al. 2012). Both the Hog1-mCh strain and the NLS-Hog1 strain restored *ENA1* induction within 60 min of NaCl exposure, whereas the Hog1-CAAX strain exhibited markedly reduced *ENA1* induction that was significantly lower than wild-type levels (*P* = 0.019) and did not significantly differ from that of the *hog1*Δ mutant (*P* = 0.11) (Fig. 2c). These findings indicate that nuclear translocation of Hog1 is essential for full transcriptional activation of *ENA1* during osmoadaptation.

Intracellular glycerol biosynthesis is another critical process for osmoadaptation. In *S. cerevisiae*, *GPD1* and *GPD2* encode glycerol-3-phosphate dehydrogenases that catalyze the reversible conversion of dihydroxyacetone phosphate to glycerol-3-phosphate. This intermediate is subsequently converted to glycerol, an essential intracellular osmoprotectant, by glycerol-3-phosphatases (*GPP1* and *GPP2*) (Lee et al. 2012; Pahlman et al. 2001). In *C. neoformans*, *GPD1* (CNAG_00121) and *GPD2* (CNAG_01745) perform analogous functions, and four *GPP1*/*GPP2* orthologs have been identified: *HAD1* (*GPP2*, CNAG_01774), *HAD6* (CNAG_06122), *HAD8* (CNAG_06132), and *HAD9* (CNAG_06698) (Martho et al. 2019). Among these, *HAD1* has previously been shown to be regulated by Hog1 (Ko et al. 2009). Under NaCl-induced osmotic stress, *GPD1* expression increased in the wild-type strain but not significantly in the *hog1*Δ mutant (Fig. 2c). Expression was restored to wild-type levels in both the Hog1-mCh and Hog1-CAAX strains, whereas the NLS-Hog1 strain displayed *GPD1* expression levels comparable to those of the *hog1*Δ mutant. In contrast, *GPD2* expression slightly increased in WT, Hog1-mCh, and Hog1-CAAX at 60 min, while the *hog1*Δ and NLS-Hog1 mutants exhibited lower basal *GPD2* induction and showed no induction of *GPD2* (Fig. 2c), suggesting that glycerol biosynthesis can be induced by cytoplasmic Hog1 function.

Among the glycerol-3-phosphatase orthologs, *HAD1* expression increased in the wild-type but remained unchanged in the *hog1*Δ mutant (Supplementary Fig. S4a). Expression was restored in the Hog1-mCh and Hog1-CAAX strains, similar to the regulation observed for *GPD1* or *GPD2*. Notably, after 60 min of treatment, *HAD1* expression was higher in Hog1-CAAX than in the wild type, suggesting hyperactivation of cytoplasmic transcriptional activators. Under osmotic stress, *HAD9* induction was abolished in *hog1*Δ. In NLS-Hog1, there was a statistically significant induction compared to the unstressed condition, whereas the induction in Hog1-CAAX did not reach statistical significance (*P* = 0.1714), although the magnitude of induction was comparable between the two strains. These results suggest that nuclear localization of Hog1 may contribute to, but is not solely responsible for, full *HAD9* induction (Supplementary Fig. S4a). However, expression levels of *HAD6* and *HAD8* remained unchanged across Hog1 deletion, suggesting that their transcriptional regulation is independent of Hog1 (Supplementary Fig. S4a). Collectively, these findings suggest that cytoplasmic and nuclear localization of Hog1 distinctly contribute to transcriptional activation of osmoadaptive effector genes in *C. neoformans* (Fig. 2d).

### Membrane-tethered Hog1 causes excessive glycerol accumulation independently of transcriptional upregulation of glycerol biosynthetic genes in *C. neoformans*

Because the highly enhanced osmotic susceptibility of the membrane-tethered Hog1 variant was unlikely to result from altered expression of osmoadaptive effector genes, we hypothesized that membrane-tethered Hog1 might more efficiently, or even aberrantly, stimulate intracellular glycerol biosynthesis compared with the wild-type and nuclear Hog1 variants. As the relevant metabolic enzymes are cytoplasmic, this could result in excessive glycerol accumulation, osmotic imbalance, and subsequent hypersensitivity.

To test this possibility, we assessed susceptibility to fludioxonil, a phenylpyrrole fungicide known to hyperactivate the fungal HOG pathway, leading to excessive glycerol overproduction and growth arrest (Bahn 2008; Kojima et al. 2006). Supporting our hypothesis, the Hog1-CAAX strain displayed markedly increased susceptibility to fludioxonil compared with the wild-type strain, whereas the NLS-Hog1 and Hog1-mCh strains retained near wild-type resistance levels (Fig. 3a). Notably, even at a low concentration of fludioxonil (0.5 μg/mL), which did not affect the wild type, the Hog1-CAAX strain exhibited marked growth inhibition (Fig. 3a).

**Fig. 3.**
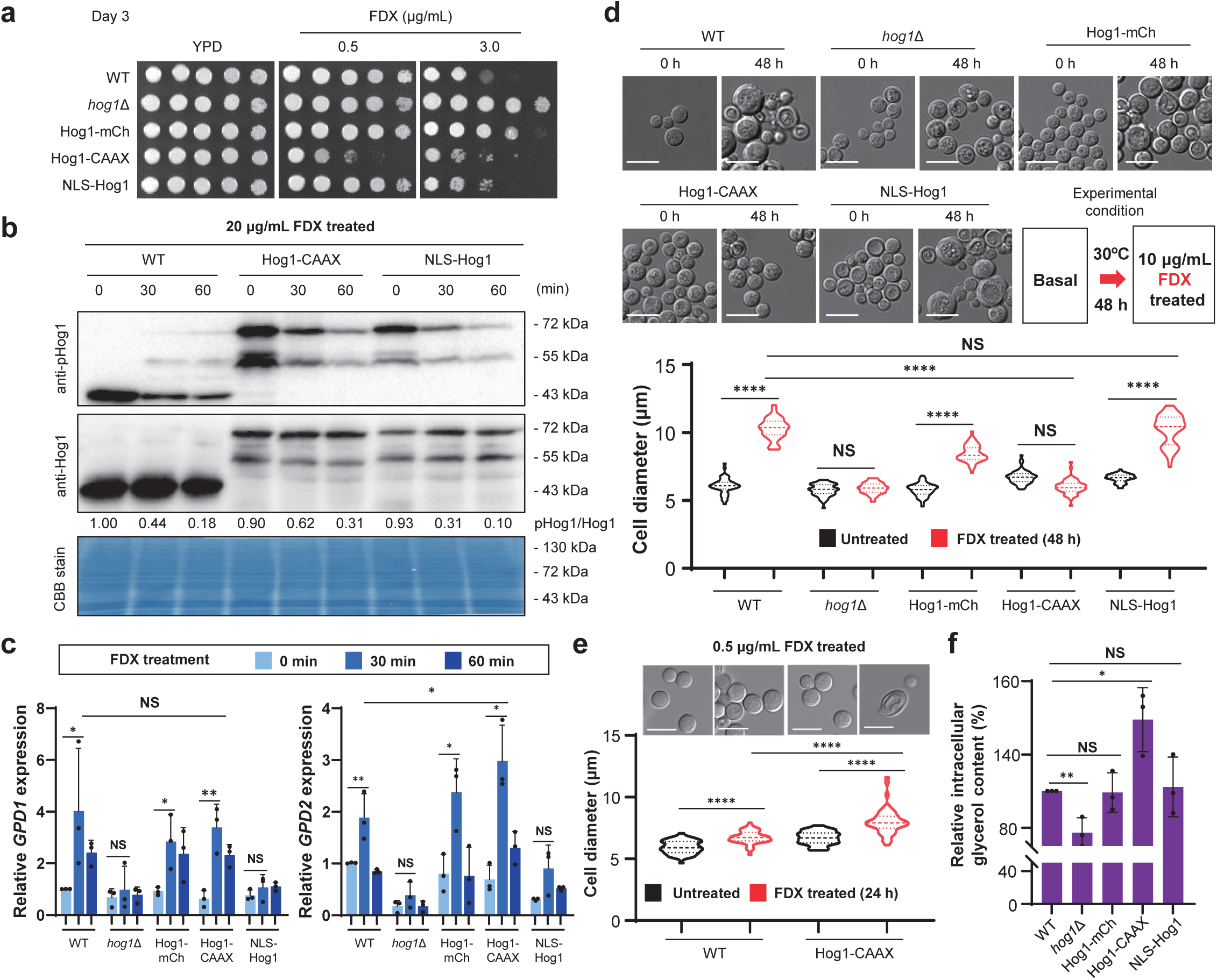
Cytoplasmic Hog1 induced excessive glycerol accumulation under fludioxonil treatment. (a) Phenotypic assay assessing susceptibility to fludioxonil at different concentrations (0.5 and 3 µg/mL). Ten-fold serial dilutions (10^-4^ to undiluted cells) were spotted and incubated for 3 days at 30°C. (b) Western blot demonstrating Hog1 phosphorylation status in response to fludioxonil treatment (20 µg/mL). Hog1 phosphorylation was detected using a phospho-p38 antibody, and total Hog1 levels were detected using an anti-Hog1 antibody. Coomassie Brilliant Blue (CBB) staining of the membrane was used as a loading control. The ∼43 kDa band in the WT lane corresponds to endogenous native Hog1, which is absent in the mCherry-tagged Hog1 strains. The ∼55 kDa band in the Hog1-mCh, Hog1-CAAX, and NLS-Hog1 lanes represents a proteolytic cleavage product of the Hog1-mCherry fusion proteins (see Fig. 1b for characterization). (c) qRT-PCR analysis of glycerol-3-phosphate dehydrogenase *GPD1* and *GPD2* expression levels following fludioxonil exposure (10 µg/mL). Expression values were normalized as described above. (d) Representative microscopic images illustrating morphological changes in wild-type (WT), *hog1*Δ, Hog1-mCh, Hog1-CAAX, and NLS-Hog1 mutants after treatment with fludioxonil (10 µg/mL for 48 h). Scale bars represent 10 µm. Cell diameter was quantified under control conditions and following fludioxonil treatment. Data points represent measurements from 50 individual cells, and horizontal bars indicate mean values. The black violin plot represents untreated cells, while the red plot corresponds to fludioxonil-treated cells, consistent with the images shown in the upper panel. **** indicates *P* < 0.0001; NS indicates a non-significant difference. (e) WT and Hog1-CAAX cells were treated with a low concentration of fludioxonil (0.5 µg/mL for 24 h), followed by microscopy imaging and quantification. Quantification was performed using the same method as described above. (f) Relative intracellular glycerol content was measured in the strains used in (Fig. 3) using a commercial glycerol assay kit.

To determine whether this hypersusceptibility of Hog1-CAAX was linked to altered Hog1 phosphorylation dynamics, we monitored Hog1 phosphorylation levels in wild-type and mutants under fludioxonil treatment. Unexpectedly, all strains, including Hog1-CAAX and NLS-Hog1, exhibited dephosphorylation patterns similar to the wild type (Fig. 3b), indicating that the observed hypersusceptibility was not attributable to altered Hog1 phosphorylation kinetics. In addition, the change in phosphorylation was slightly delayed in Hog1-CAAX, consistent with the response to osmotic stress in *C. neoformans* (Fig. 2b; Supplementary Fig. S3b).

We next examined the expression of glycerol biosynthesis genes (*GPD1/2* and *HAD1/6/8/9*) under fludioxonil treatment. As previously reported, *GPD1/2* induction was strongly reduced in the *hog1*Δ mutant. Interestingly, the Hog1-CAAX strain restored *GPD1/2* expression to wild-type levels, whereas the NLS-Hog1 strain did not show statistically significant induction of either *GPD1* or *GPD2* (Fig. 3c). This finding further suggests that the cytoplasmic Hog1 may activate transcription of *GPD1/2* indirectly, potentially via a cytoplasmic transcription factor(s). In contrast, none of *HAD1/6/8/9* genes were induced under these conditions, indicating that their expression is not regulated by Hog1 under fludioxonil exposure (Supplementary Fig. S4b). Collectively, these results indicate that the fludioxonil hypersusceptibility of the Hog1-CAAX strain is not due to transcriptional dysregulation of glycerol biosynthetic genes.

We further analyzed cell morphology after fludioxonil treatment. Following 48 h of exposure to a high concentration (10 µg/mL), wild-type cells exhibited marked swelling, whereas the *hog1*Δ mutant did not, consistent with previous reports (Kim et al. 2011). Unexpectedly, the Hog1-CAAX strain also failed to swell under these conditions (Fig. 3d). This absence of enlargement likely reflects immediate and severe osmotic imbalance, resulting in transient cell swelling followed by cell lysis. To explore this possibility, we treated the Hog1-CAAX mutant with a low fludioxonil concentration (0.5 µg/mL) and monitored for morphological changes. After 24 h, Hog1-CAAX cells displayed significantly greater swelling than wild-type cells (Fig. 3e), suggesting accelerated and excessive intracellular glycerol accumulation. To directly assess this, we quantified intracellular glycerol content (Fig. 3f). Even under unstressed conditions, the Hog1-CAAX strain contained higher basal glycerol levels than the wild type, whereas the *hog1*Δ mutant had reduced levels.

Taken together, these findings demonstrate that the membrane-tethered Hog1 enhances glycerol biosynthesis independently of transcriptional upregulation of glycerol biosynthetic genes in *C. neoformans*. These data suggest that cytoplasmic or membrane-associated Hog1 modulates glycerol metabolism at the post-translational level, for example by influencing the activity of metabolic enzymes or glycerol transport, thereby contributing to excessive glycerol accumulation and osmotic imbalance in the fungal pathogen.

### Both membrane-tethered and nuclear Hog1 are required for thermal, oxidative, and membrane stress responses in *C. neoformans*

To further elucidate how Hog1 subcellular localization influences stress adaptations, we examined the stress tolerance of Hog1 localization variants under diverse environmental challenges. Hog1 is essential for thermotolerance in *C. neoformans*, particularly at elevated temperatures (e.g., 39-40°C). While the Hog1-CAAX mutant partially restored thermotolerance, the NLS-Hog1 strain fully rescued the growth defect at high temperature (Fig. 4a). These results indicate that both membrane-tethered and nuclear Hog1 contribute to thermotolerance.

**Fig. 4.**
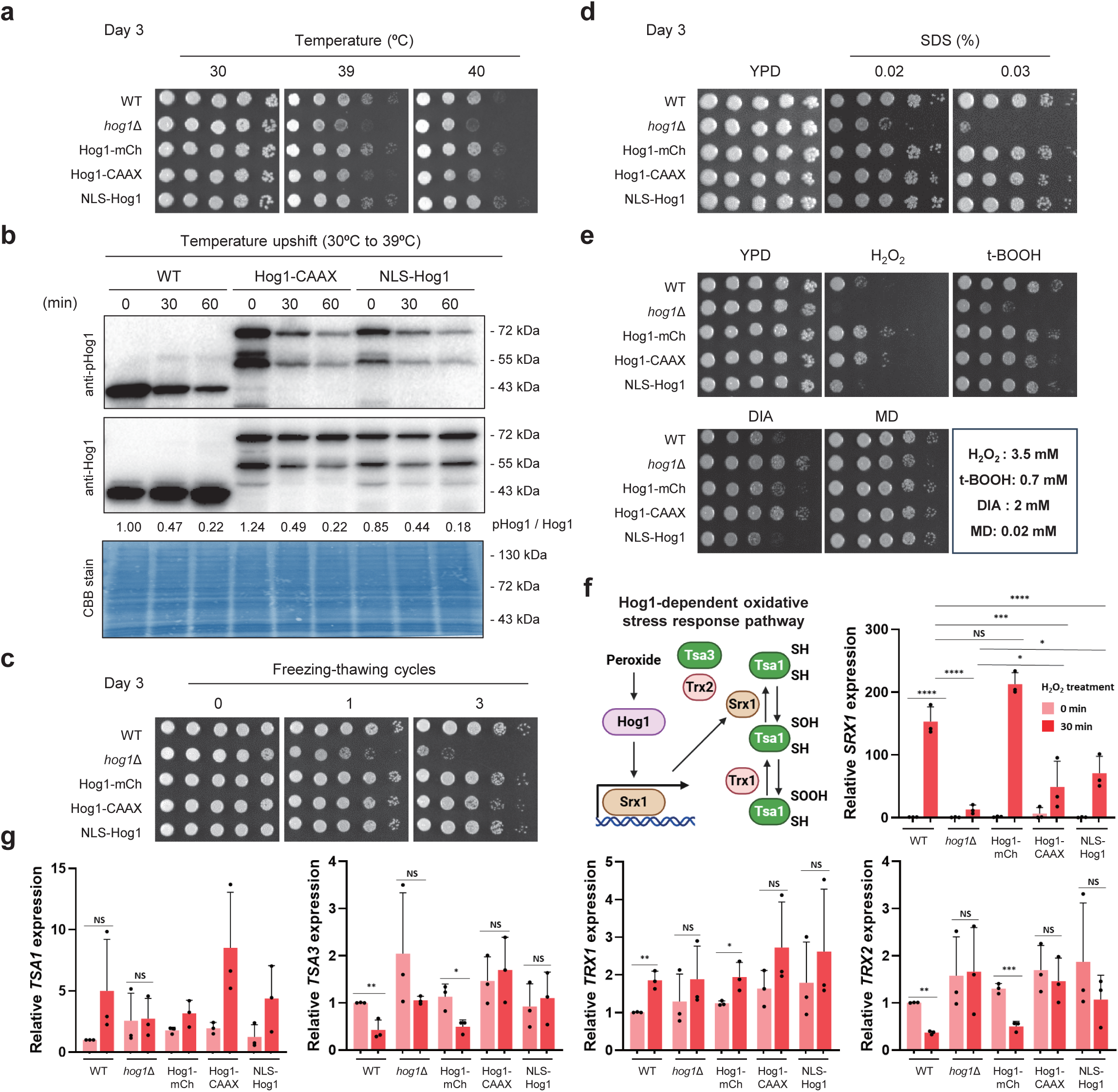
Cytoplasmic or nuclear Hog1 is dispensable for membrane stability of *C. neoformans*. (a) Wild-type (WT), *hog1*Δ, Hog1-mCh, Hog1-CAAX, and NLS-Hog1 mutants were spotted onto YPD medium and incubated for 3 days at various temperatures, after which images were taken. (b) WT, Hog1-CAAX, and NLS-Hog1 were synchronized to the same growth phase, then subjected to a temperature upshift from 30°C to 39°C. Proteins were extracted at 0 (basal, pre-stress), 30, and 60 min following the upshift. Hog1 phosphorylation was detected using a phospho-p38 antibody, and total Hog1 levels were detected using an anti-Hog1 antibody. Coomassie Brilliant Blue (CBB) staining of the membrane was used as a loading control. The ∼43 kDa band in the WT lane corresponds to endogenous native Hog1, which is absent in the Hog1-CAAX and NLS-Hog1 strains. The ∼55 kDa band in the Hog1-CAAX and NLS-Hog1 lanes represents a proteolytic cleavage product of the Hog1-mCherry fusion proteins (see Fig. 1b for characterization). (c-e) To assess freezing-thawing resistance using the same strains as in Fig 4a, cells were subjected to 0, 1, and 3 cycles of freezing in liquid nitrogen followed by thawing, then spotted onto YPD medium. Growth was examined after 3 days of incubation. Additionally, to evaluate stress resistance, cells were spotted onto YPD medium supplemented with the membrane stressor sodium dodecyl sulfate (SDS) and the oxidative stressors hydrogen peroxide (H_2_O_2_), *tert*-butyl hydroperoxide (t-BOOH), diamide (DIA), and menadione (MD). The growth was monitored after 3 days. (f-g) A schematic diagram of the known Hog1 involved in sulfiredoxin regulation is shown (https://BioRender.com/tkokm2e). Using the strains from Fig. 4a, the expression levels of the genes *SRX1*, *TSA1*, *TSA3*, *TRX1*, and *TRX2*, which are associated oxidative stress response, were analyzed before and after hydrogen peroxide treatment using qRT-PCR. Statistical analysis was performed using ANOVA; **** indicates *P* < 0.0001, *** indicates *P* < 0.001, ** indicates *P* < 0.01, * indicates *P* < 0.05, and NS indicates no statistically significant difference.

To determine whether this phenotype correlated with altered Hog1 activation, we analyzed Hog1 phosphorylation following a temperature shift from 30°C to 39°C. While all three strains underwent dephosphorylation upon heat exposure, NLS-Hog1 reached lower phospho-Hog1/Hog1 ratios than Hog1-CAAX by 60 min (Fig. 4b; Supplementary Fig. S3d), a pattern consistent with the more efficient dephosphorylation of nuclear Hog1 also observed under osmotic stress (Fig. 2b; Supplementary Fig. S3b) and fludioxonil treatment (Fig. 3b; Supplementary Fig. S3c). Notably, the wild-type and Hog1-CAAX strains showed comparable phospho-Hog1/Hog1 ratios at 60 min in the optimized experiment, indicating that the mild thermosensitivity of the Hog1-CAAX strain is unlikely to result primarily from differences in Hog1 phosphorylation dynamics.

Hog1 also mediates cold stress resistance. Previous studies showed that *hog1*Δ mutants are highly susceptible to freeze-thaw cycles (So et al. 2020). The Hog1-mCh and NLS-Hog1 strains fully restored cryotolerance to wild-type levels, whereas the Hog1-CAAX strain, though less sensitive than *hog1*Δ, exhibited slightly reduced survival compared to wild type (Fig. 4c). These findings suggest that while membrane-tethered Hog1 contributes to cryostress adaptation, nuclear Hog1 alone is sufficient for survival. Given that temperature fluctuations can disrupt membrane integrity and fluidity, we also tested sensitivity to the membrane destabilizer sodium dodecyl sulfate (SDS). Notably, both membrane-tethered and nuclear Hog1 restored wild-type resistance to SDS (Fig. 4d).

Thermal stress often increases reactive oxygen species (ROS) production by accelerating cell metabolism, thereby imposing oxidative stress (Musa et al. 2018; Slimen et al. 2014). To assess whether membrane-tethered and nuclear Hog1 pools are both required for oxidative stress resistance, we tested growth under various ROS-generating agents, including hydrogen peroxide (H_2_O_2_), organic peroxide (*tert*-butyl hydroperoxide, t-BOOH), thiol oxidizing reagent (diamide, DIA), superoxide generator (menadione, MD) (Fig. 4e). Consistent with previous reports, the *hog1*Δ mutant was hypersensitive to H_2_O_2_, t-BOOH, and MD, but resistant to DIA. These stress phenotypes were largely rescued in the Hog1-mCh strain. The NLS-Hog1 strain exhibited a similar phenotypic rescue pattern except under H_2_O_2_ stress (Fig. 4e), where it remained moderately sensitive. Interestingly, the Hog1-CAAX strain restored wild-type resistance to H_2_O_2_, t-BOOH, and MD, but not to DIA (Fig. 4e), suggesting that membrane-tethered Hog1 can activate cytoplasmic ROS detoxification pathways.

To further substantiate this, we analyzed the expression of cytoplasmic oxidative stress response genes, including *SRX1, TSA1/3,* and *TRX1/2* (Upadhya et al. 2013). As previously reported, *SRX1* expression was induced over 100-fold in wild-type cells under oxidative stress but abolished in the *hog1*Δ mutant. This induction was fully restored in the Hog1-mCh strain but only partially recovered in both Hog1-CAAX and NLS-Hog1 mutants (Fig. 4f). Under oxidative conditions, *TRX1* expression increased in wild-type and Hog1-mCh strains, while *TSA3* and *TRX2* expression decreased. In contrast, expression of *TRX1*, *TRX2*, and *TSA3* remained largely unchanged in *hog1*Δ, Hog1-CAAX, or NLS-Hog1 mutants. *TSA1* expression was unaffected across all strains (Fig. 4g).

Collectively, these findings indicate that the Hog1 subcellular localization differentially regulates transcriptional response to oxidative stress, with both cytoplasmic and nuclear pools contributing to full oxidative defense functionality in *C. neoformans*.

### Distinct roles of membrane-tethered and nuclear Hog1 in ER stress response

Hog1 plays a crucial role in the cellular adaptation to ER stress. Consistent with previous findings, the *hog1*Δ mutant was highly susceptible to tunicamycin (TM), an inducer of ER stress (Cheon et al. 2014). To determine how Hog1 localization influences the ER stress response, we examined the TM susceptibility of the Hog1-CAAX and NLS-Hog1 mutants. The NLS-Hog1 strain fully restored TM resistance to wild-type levels, whereas the Hog1-CAAX strain exhibited pronounced hypersensitivity (Fig. 5a). These results indicate that the nuclear activity of Hog1 is essential for survival under ER stress.

**Fig. 5.**
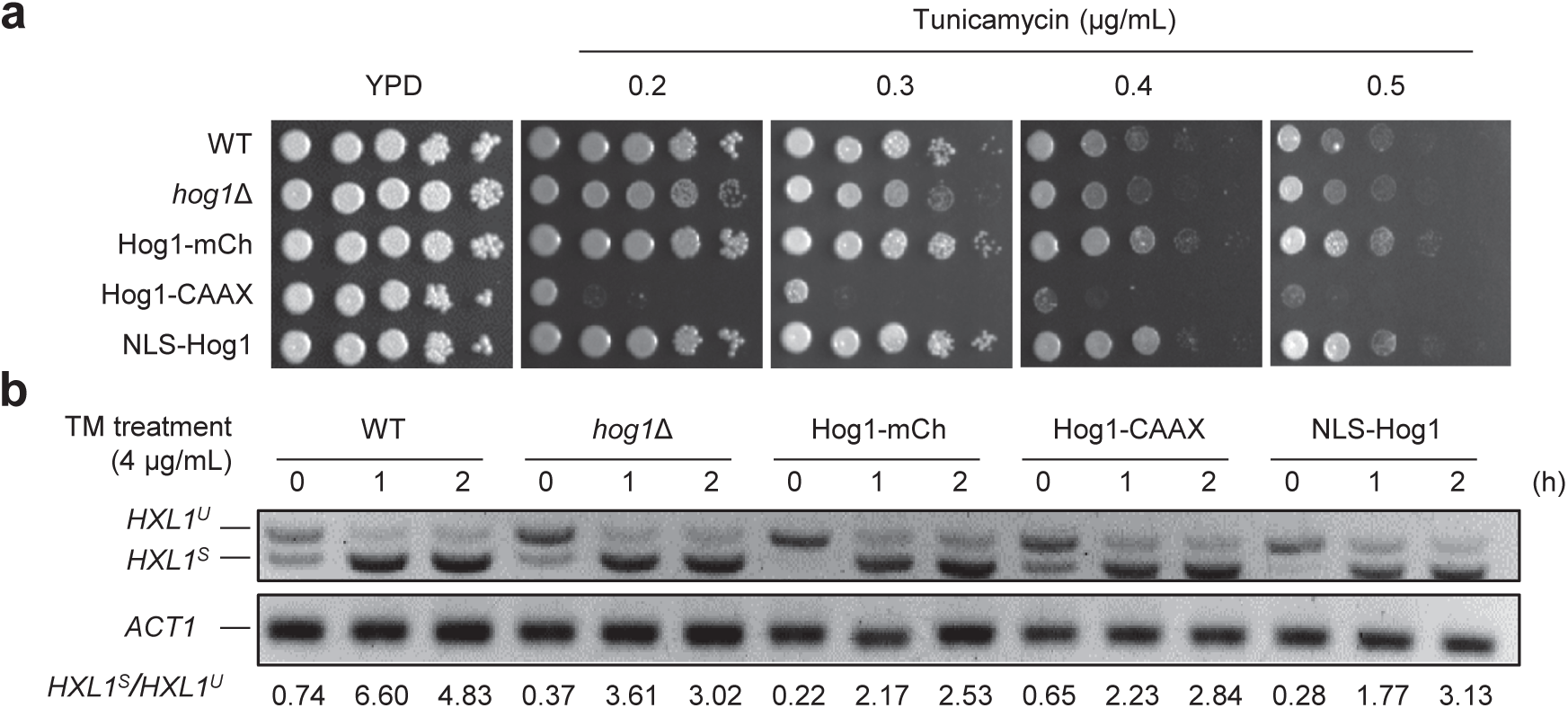
Membrane-tethered Hog1 regulates ER stress response with noncanonical pathway. (a) Phenotypic assay of wild-type (WT), *hog1*Δ, Hog1-mCh, Hog1-CAAX, and NLS-Hog1 mutants on YPD agar supplemented with various concentrations of tunicamycin (0.2 to 0.5 µg/mL) and further incubated 3 days. (b) To visualize and quantify *HXL1* splicing, WT and the previously used Hog1 mutants were grown to saturation over two days, then treated with 4 µg/mL tunicamycin for 0, 1, and 2 h. Total RNA was extracted, followed by cDNA synthesis, and spliced and unspliced forms of *HXL1* were visualized by RT-PCR amplification. Housekeeping gene *ACT1* was used as a control gene.

The extreme hypersensitivity observed in the Hog1-CAAX strain suggests that mislocalized, membrane-tethered Hog1 not only fails to confer protection but may even exacerbate cellular dysfunction under ER stress, possibly due to inappropriate or excessive signaling. However, the splicing of *HXL1* mRNA, a hallmark of UPR activation mediated by the Ire1-Hxl1 pathway was comparable across all strains (Fig. 5b). This finding demonstrates that the protective function of nuclear Hog1 under ER stress operates independently of the canonical Ire1-Hxl1 UPR signaling cascade. Instead, Hog1 may promote ER stress resistance via a parallel, non-canonical pathway or through regulation of other proteostasis-related genes independently of Hxl1-driven transcription. Alternatively, nuclear Hog1 may enhance compensatory mechanisms that maintain protein homeostasis under ER stress conditions.

### Distinct roles of membrane-tethered and nuclear Hog1 in ergosterol biosynthesis

Hog1 has also been shown to exert opposing effects on resistance to polyene and azole antifungal agents. Amphotericin B (AMB), a polyene, binds to ergosterol in the fungal cell membrane, forming pores that compromise membrane integrity and cause cell death. In contrast, azole drugs such as fluconazole (FLC), voriconazole (VRC), and ketoconazole (KTC) inhibit lanosterol 14a-demethylase, a key enzyme in ergosterol biosynthesis, thereby impairing membrane structure and function. Previous studies demonstrated that *HOG1* deletion increases AMB susceptibility while paradoxically enhancing azole resistance in *C. neoformans* (Kim et al. 2011; Ko et al. 2009), implicating the HOG pathway in the regulation of ergosterol biosynthesis.

In this study, we found that both Hog1-CAAX and NLS-Hog1 strains fully restored resistance to AMB (Fig. 6a), suggesting that cytoplasmic Hog1 activity is sufficient to mediate AMB resistance. Upon azole treatments (FLC, VRC, and KTC), however, the NLS-Hog1 strain exhibited resistance comparable to the *hog1*Δ mutant, while the Hog1-CAAX strain not only reinstated azole susceptibility but displayed even greater susceptibility than the wild type (Fig. 6a).

**Fig. 6.**
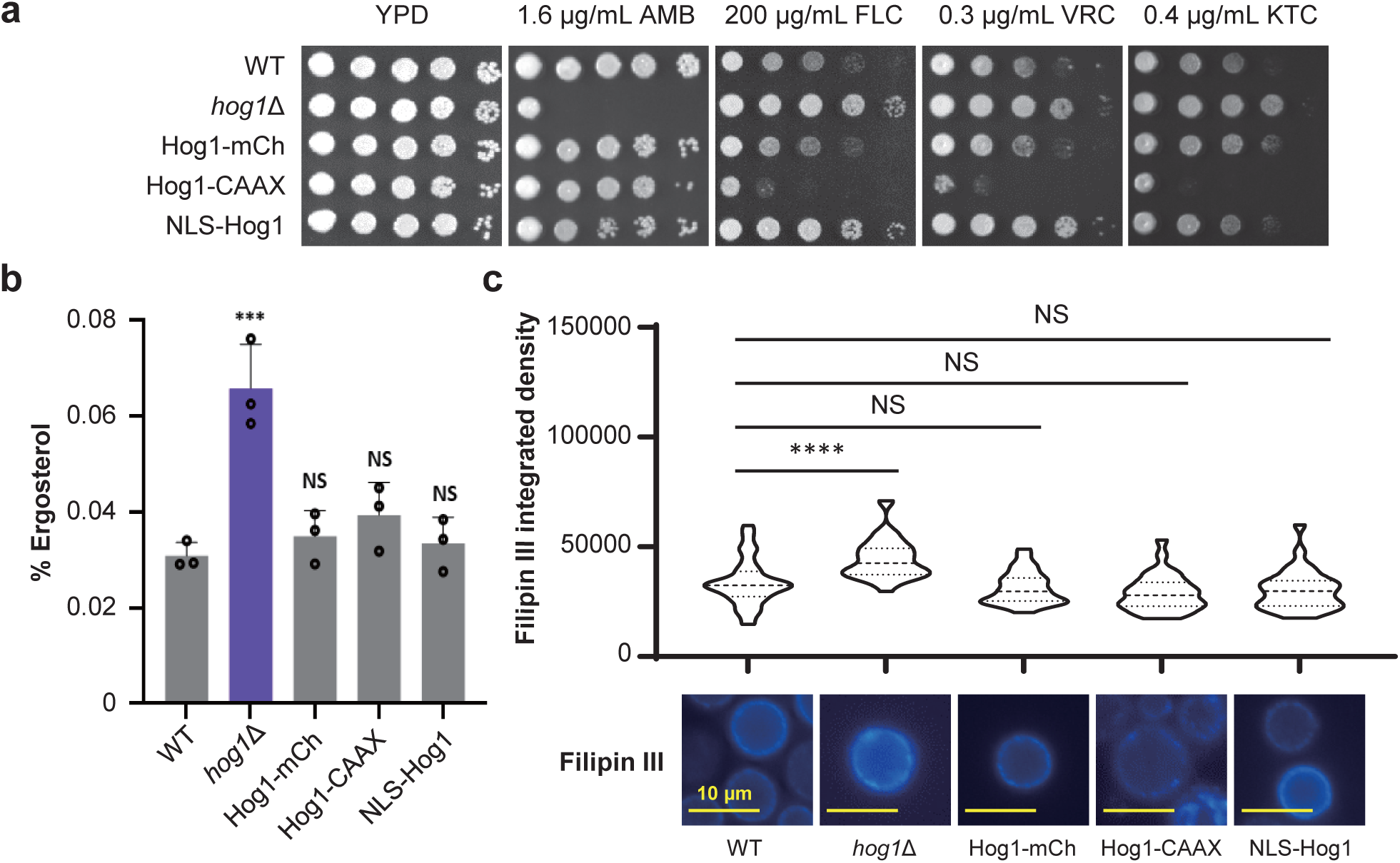
Hog1 regulates basal ergosterol synthesis regardless of subcellular location. (a) Phenotypic assay of wild-type, *hog1*Δ, Hog1-mCh, Hog1-CAAX, and NLS-Hog1 mutants on YPD supplemented with indicated concentration of amphotericin B (AMB), fluconazole (FLC), voriconazole (VRC), or ketoconazole (KTC). Plates were further incubated with 30°C after 3 days. (a) Ergosterol content of the indicated strains was measured with the ergosterol assay (described in Materials and Methods). *** indicates *P* < 0.001; NS indicates a non-significant difference. (c) Ergosterol of the previously described strains was stained with Filipin III, and their fluorescence intensity was measured by ImageJ. **** indicates *P* < 0.0001; NS indicates a non-significant difference. The scale bar represents 10 µm.

To determine whether these phenotypes correlated with changes in ergosterol content, we quantified ergosterol levels using a biochemical ergosterol assay. As expected, the *hog1*Δ mutant accumulated elevated ergosterol levels, and reintroduction of Hog1-mCh restored ergosterol content to wild-type levels, irrespective of its subcellular localization (Fig. 6b). Filipin III staining, which specifically binds ergosterol, confirmed these results (Fig. 6c).

Together, these results indicate that both cytoplasmic and nuclear Hog1 contribute to maintaining proper ergosterol homeostasis. Cytoplasmic Hog1 primarily supports resistance to polyene-induced membrane damage, whereas nuclear Hog1 may participate in transcriptional regulation of ergosterol biosynthetic or stress-responsive genes. Thus, the coordinated activities of both Hog1 pools are critical for preserving membrane integrity under antifungal stress conditions.

### Both membrane-tethered and nuclear Hog1 are critical for regulating mating

In *C. neoformans*, Hog1 functions as a key negative regulator of the pheromone-responsive MAPK cascade. Deletion of *HOG1* leads to hyperactivation of sexual differentiation, characterized by elevated expression of the mating pheromone gene *MFa1*, enhanced cell fusion efficiency, and thereby extensive filamentation during mating (Bahn et al. 2005). In this study, we found that both the Hog1-mCh and Hog1-CAAX strains exhibited filamentation levels comparable to wild type, indicating that membrane-tethered Hog1 is sufficient to repress sexual differentiation. In contrast, the NLS-Hog1 strain still displayed pronounced filamentation, although to a lesser extent than the *hog1*Δ mutant, demonstrating that nuclear Hog1 alone is insufficient to fully suppress mating (Fig. 7a). Despite these differences in filamentation, all Hog1 localization variants were capable of forming mating filaments, producing basidia, and completing sporulation (Fig. 7b). These observations suggest that Hog1 subcellular localization primarily affects the initiation of sexual development rather than the later morphogenetic processes of filamentation and sporulation.

**Fig. 7.**
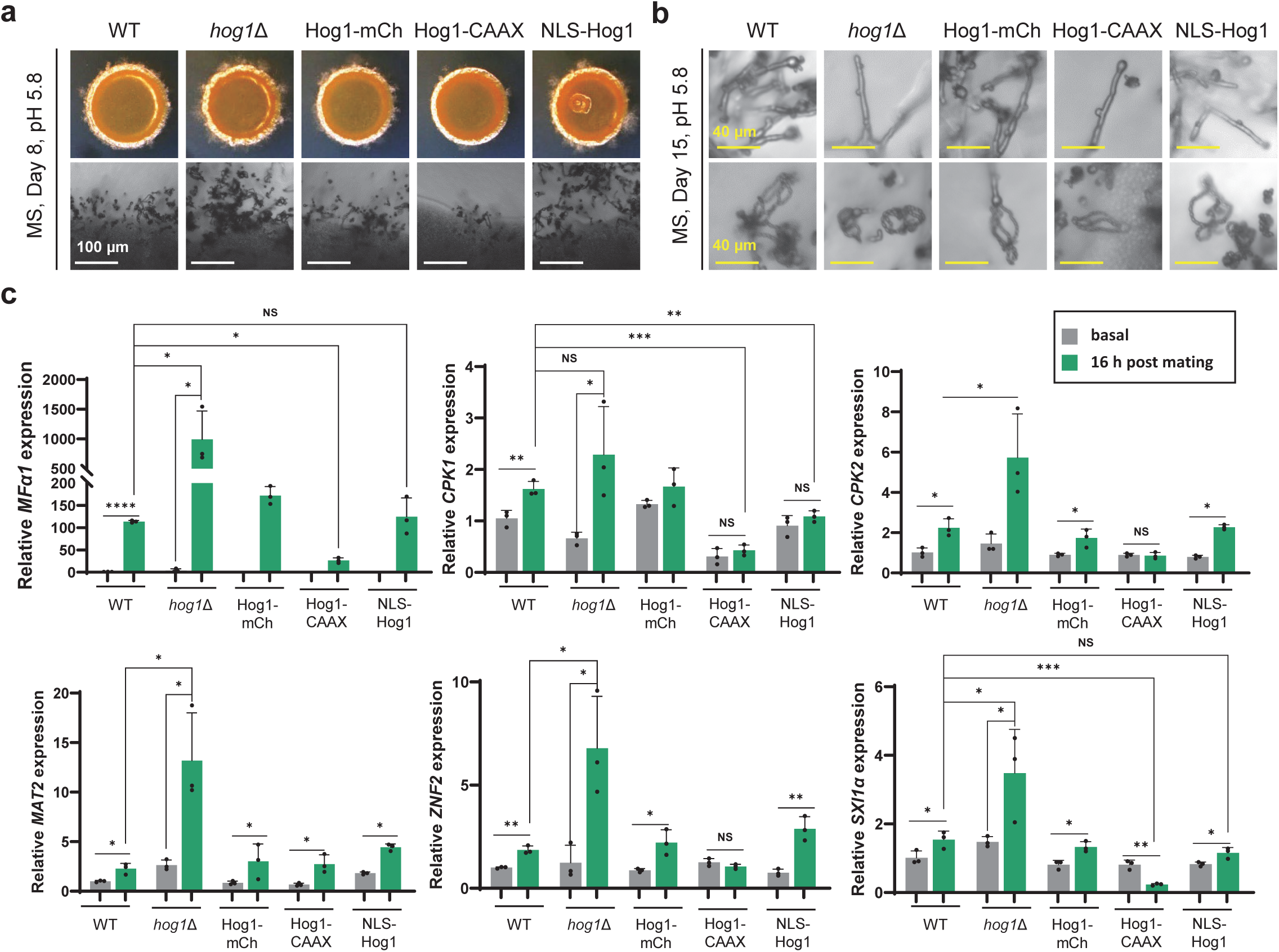
Cytoplasmic and nuclear Hog1 regulate sexual development. (a, b) *MAT*a wild-type H99, *hog1*Δ, Hog1-mCh, Hog1-CAAX, and NLS-Hog1 mutants were mating with *MAT***a** wild-type YL99 at pH 5.8 MS medium (Murashige & Skoog medium) and grown at room temperature in the dark for 8 days (a) or 15 days (b). Colonies, filaments, and spore chains were then observed under a brightfield microscope. White scale bar indicates 100 µm and yellow scale bar indicates 40 µm. (c) The same strain as in Fig. 7a was used. Equal amounts of cultured cells were spread onto V8 medium and harvested both prior to mating and after 16 h of incubation under dark conditions at room temperature. The transcription level of mating-related genes (*MFa1, CPK1, CPK2, MAT2, ZNF2*, and *SXI1a*) were quantified with qRT-PCR. Statistical analysis was performed using Student’s *t*-test. Significance levels are indicated as follows: *, *P* < 0.05; **, *P* < 0.01; ***, *P* < 0.001; ****, *P* < 0.0001; NS, not significant.

To further substantiate these findings, we examined the expression of key mating-responsive genes, including *MFa1, CPK1, CPK2, MAT2, ZNF2*, and *SXI1a*, in the Hog1 localization mutants (Jang et al. 2024). Consistent with its hyperfilamentation phenotype, the *hog1*Δ mutant exhibited significantly elevated transcript levels of all tested genes following mating (Fig. 7c). In contrast, the Hog1-CAAX strain showed reduced expression of these genes, confirming that membrane-tethered Hog1 effectively represses the Cpk1 MAPK pathway. Interestingly, the NLS-Hog1 strain, despite its enhanced filamentation, exhibited near wild-type expression levels for most mating-related genes.

Collectively, these findings indicate that both cytoplasmic and nuclear Hog1 contribute to the regulation of mating and associated gene expression in *C. neoformans*. However, cytoplasmic Hog1 plays a prominent role in repressing pheromone signaling and preventing excessive sexual differentiation.

### Both membrane-tethered and nuclear Hog1 distinctly regulate virulence factor production

Beyond its functions in stress adaptation and sexual differentiation, Hog1 also acts as a negative regulator of three major virulence factors in *C. neoformans*: the polysaccharide capsule, polyphenol melanin pigments, and urease (Bahn et al. 2005; Lee et al. 2016). The capsule, which forms the outermost layer of the fungal cell wall, protects *C. neoformans* from phagocytosis by host macrophages (Jang et al. 2022). Melanin, anchored to the inner cell wall through chitin, serves as both an antioxidant and an antiphagocytic factor (Lee et al. 2019). Urease facilitates traversal of the blood-brain barrier, thereby promoting fungal dissemination and central nervous system invasion (Singh et al. 2013).

To evaluate the impact of Hog1 subcellular localization on virulence factor regulation, we analyzed capsule, melanin, and urease production in the membrane-tethered and nuclear Hog1 mutants. Both the Hog1-mCh and NLS-Hog1 strains restored capsule production to wild-type levels, whereas the Hog1-CAAX strain exhibited an intermediate phenotype between wild type and the *hog1*Δ mutant (Fig. 8a). These results suggest that while both cytoplasmic and nuclear Hog1 contribute to capsule repression, nuclear localization is essential for complete suppression of capsule expansion.

**Fig. 8.**
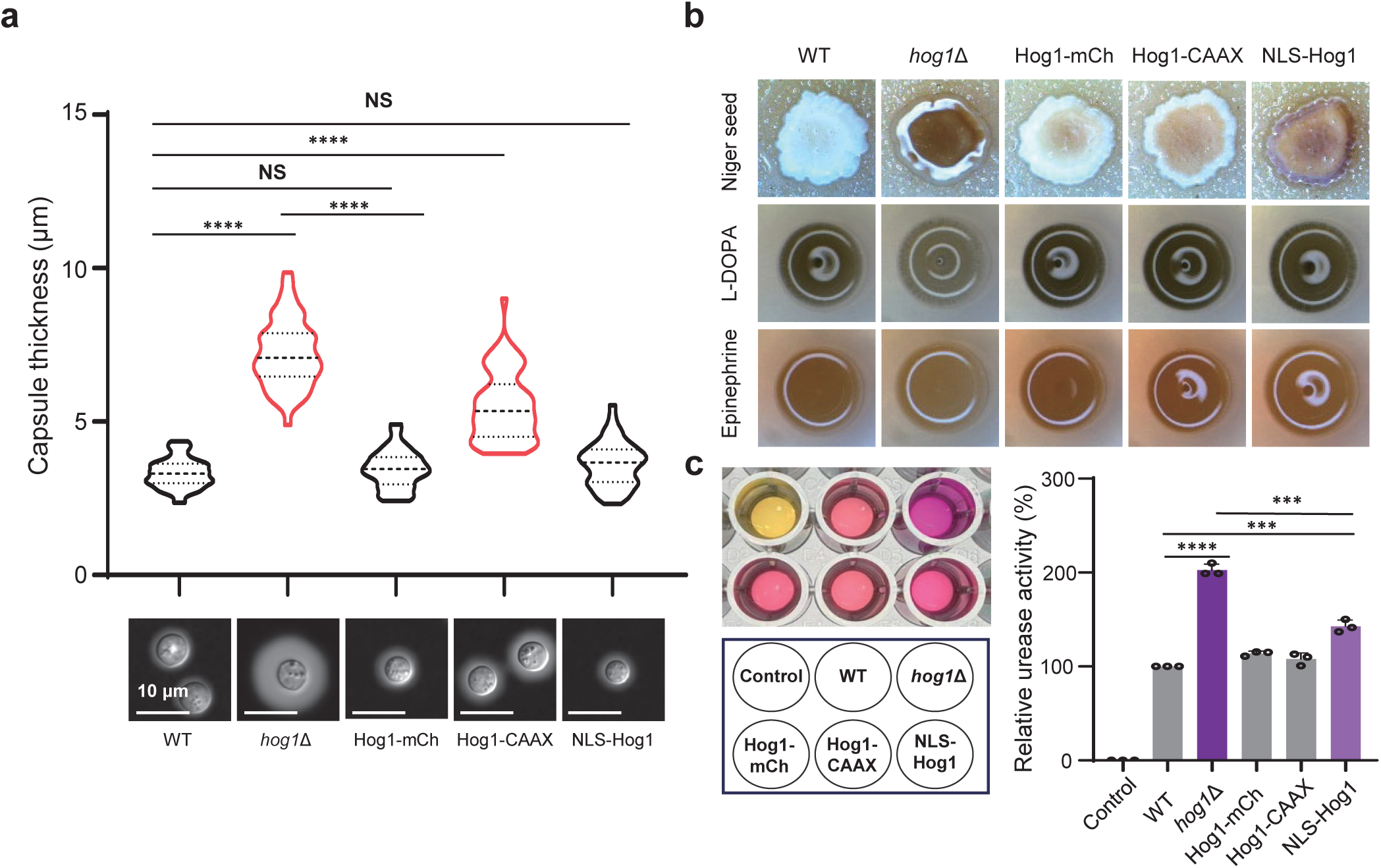
Cytoplasmic and nuclear Hog1 regulate the induction of virulence factors. (a) Wild-type (WT), *hog1*Δ, Hog1-mCh, Hog1-CAAX, and NLS-Hog1 mutants were cultured in capsule-inducing medium, DME medium (Dulbecco’s Modified Eagle Medium), at 37°C for 2 days. After India ink staining, the cells were observed under a microscope. Capsule sizes of 50 randomly selected cells were measured for quantification, and statistical analysis was performed using ANOVA. **** denotes *P* < 0.0001; NS denotes no statistically significant difference. (b) The indicated strains were spotted onto Niger seed, L-DOPA, and epinephrine medium containing 0.1% glucose, incubated at 37°C for 2 days, and then observed under a microscope. (c) An equal number of cells (10^8^) from the previously used strains were added to RUH (Rapid Urea Hydrolysis) medium and incubated at 37°C for 1 h. The absorbance at 560 nm was then measured. As a negative control, 1x RUH medium without any cells was used. Statistical analysis was performed using ANOVA; **** indicates *P* < 0.0001 and *** indicates *P* < 0.001.

For melanin production, the Hog1-mCh and Hog1-CAAX strains displayed wild-type pigmentation levels in L-DOPA or epinephrine agar (Fig. 8b). However, on Niger seed agar, the NLS-Hog1 mutant failed to fully suppress the hypermelanization phenotype observed in the *hog1*Δ mutant, indicating that nuclear Hog1 plays only a partial role in melanin regulation under specific environmental conditions. Similarly, urease activity was restored to wild-type levels in the Hog1-mCh and Hog1-CAAX strains, whereas the NLS-Hog1 mutant exhibited approximately 120% of the wild-type urease activity, reflecting incomplete repression (Fig. 8c).

Collectively, these findings suggest that Hog1 subcellular localization differentially influences the regulation of virulence-associated traits in *C. neoformans*. Both cytoplasmic and nuclear Hog1 pools contribute to controlling capsule, melanin, and urease production, yet their relative importance varies among individual virulence factors, highlighting distinct but complementary regulatory roles in fungal pathogenicity.

## Discussion

Hog1 is a central MAPK governing stress responses, developmental programs, and pathogenicity in *C. neoformans* (Bahn et al. 2005). Although this kinase dynamically shuttles between the cytoplasm and nucleus in response to external cues, the functional consequences of its compartmentalization have remained unknown. Here, by generating *C. neoformans* strains expressing plasma membrane-tethered or nuclear-restricted Hog1, we demonstrate that cytoplasmic and nuclear Hog1 make both redundant and distinct contributions to stress adaptation, sexual differentiation, ergosterol biosynthesis, and virulence factor production. The distinct and overlapping sets of effector genes influenced by Hog1 localization suggest that different subsets of transcriptional factors and regulators are differentially modulated by cytoplasmic versus nuclear Hog1, likely via phosphorylation, allowing Hog1 to coordinate diverse effectors to maintain cellular homeostasis across environmental and host niches (summarized in Fig. 9).

**Figure 9.**
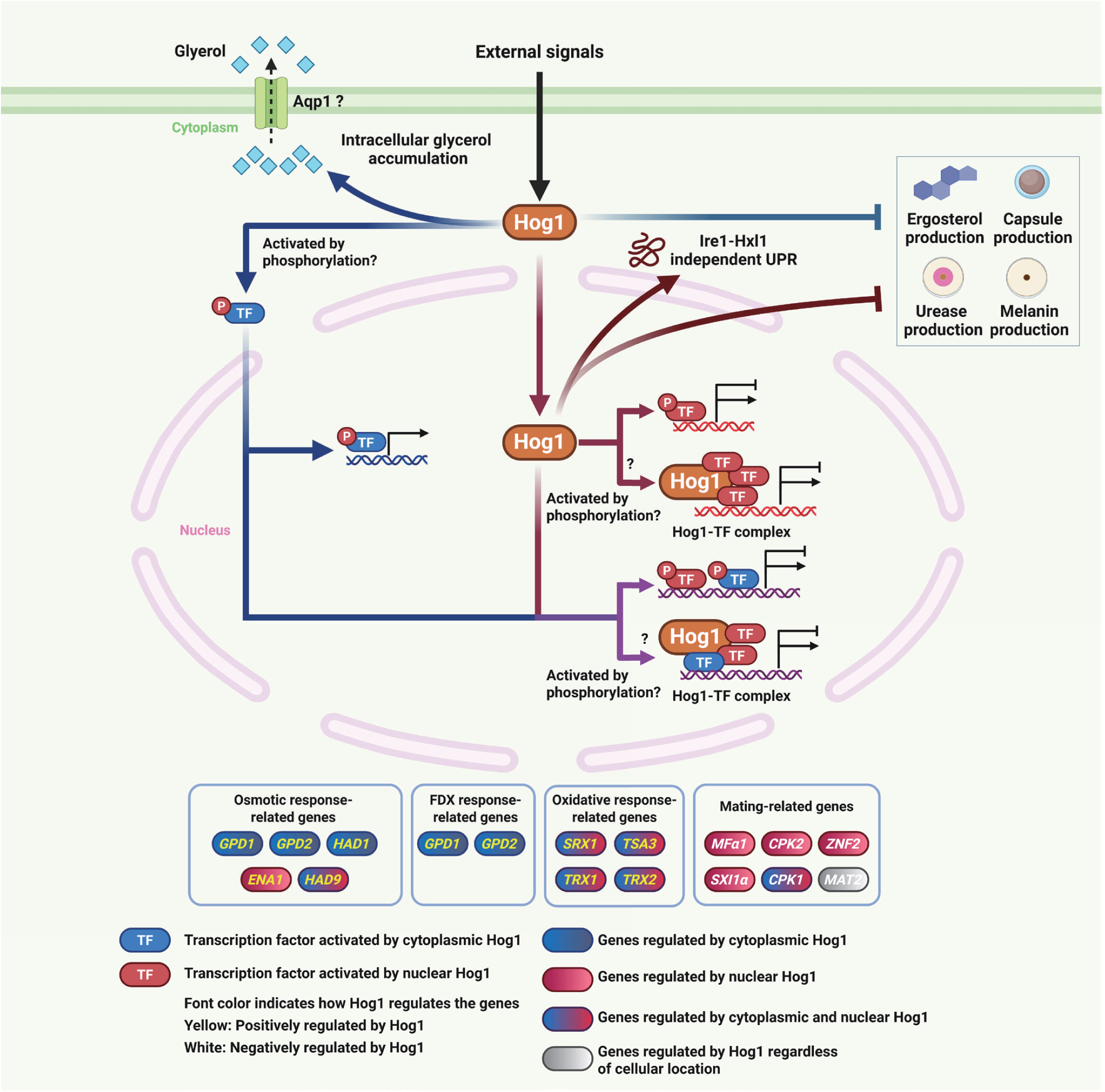
Schematic representation of subcellular localization-dependent roles of Hog1 in stress responses and pathogenic traits in *C. neoformans*. Under external stress signals, activated Hog1 exerts distinct functions depending on its subcellular localization. Cytoplasmic Hog1 promotes intracellular glycerol accumulation, potentially through post-translational regulation of the putative glycerol channel Aqp1 and activates cytoplasmic transcription factors that regulate osmotic response genes (*GPD1, GPD2, HAD1*) and fludioxonil (FDX) response genes (*GPD1*, *GPD2*). Nuclear Hog1 directs Ire1-Hxl1-independent unfolded protein response (UPR) and activates nuclear transcription factors that regulate osmotic response genes (*ENA1*), oxidative stress response genes (*SRX1*, *TSA3*, *TRX1*, *TRX2*), and mating-related genes (*MFa1*, *CPK1*, *CPK2*, *ZNF2*, *SXI1a*). Cytoplasmic and nuclear Hog1 suppress ergosterol, capsule, urease, and melanin production, and both Hog1-phosphorylated transcription factors regulate distinct, non-overlapping sets of genes under osmotic stress and oxidative stress. The mating-regulating transcription factor *MAT2* is also regulated by Hog1, but independently of its nuclear translocation. Gene box colors indicate the regulatory mode: blue, genes regulated by cytoplasmic Hog1; red, genes regulated by nuclear Hog1; blue-red gradient, genes regulated by both cytoplasmic and nuclear Hog1; gray, genes regulated by Hog1 regardless of cellular location. Gene name font colors indicate the direction of regulation by Hog1: yellow, genes positively regulated by Hog1 (stress-induced genes whose induction is abolished in the *hog1*Δ mutant); white, genes negatively regulated by Hog1 (genes whose expression is enhanced in the *hog1*Δ mutant, such as mating-related genes). Created with BioRender (https://BioRender.com/v91v944).

Our study reveals that the localization-dependent roles of Hog1 in the basidiomycete *C. neoformans* are distinct from those in the ascomycetes *S. cerevisiae* and *C. albicans* (Day et al. 2017; Westfall et al. 2008). Because all three studies employed the same Ras CAAX motif and the SV40 NLS for Hog1 targeting, phenotypic differences most likely reflect species-specific biology rather than experimental design. The most striking divergence concerns osmoresistance: in both ascomycetous yeasts, Hog1 restores osmoresistance irrespective of localization, whereas in *C. neoformans*, nuclear Hog1 fully restores wild-type resistance while membrane-tethered Hog1 further exacerbates osmosensitivity. Unexpectedly, membrane-tethered Hog1 was sufficient to induce *GPD1* and *GPD2*, whereas nuclear Hog1 was not—opposite to the pattern in *S. cerevisiae*, where nuclear Hog1 is essential for *GPD1* induction (Westfall et al. 2008), and distinct from *C. albicans*, where *GPD2* induction is Hog1 localization-independent (Day et al. 2017). These findings suggest that *GPD1/GPD2* induction in *C. neoformans* is driven by Hog1-activated cytosolic transcription factor but is not essential for osmoresistance recovery. Moreover, both localization variants exhibited wild-type-like dephosphorylation upon stress, excluding defective Hog1 phosphorylation dynamics as the cause of altered osmosensitivity.

Because the observed transcriptional and phosphorylation profiles do not account for the hypersensitivity of cells expressing membrane-tethered Hog1, we propose that sustained cytosolic Hog1 activity indirectly drives hyper-accumulation of intracellular glycerol through post-translational regulation of metabolic targets. In *S. cerevisiae*, Hog1 restricts glycerol efflux by phosphorylating the aquaglyceroporin Fps1 and inhibiting its positive regulators Rgc1/Rgc2 (Beese et al. 2009), while also activating Pfk2 (6-phosphofructo-2-kinase) to redirect glycolytic flux toward glycerol production (Dihazi et al. 2004). However, *C. neoformans* lacks clear Fps1, Rgc1, or Rgc2 orthologs—Aqp1 (CNAG_01742) is the closest Fps1 homolog, yet the *aqp1*Δ mutant is not osmosensitive (Meyers et al. 2017)—suggesting that glycerol efflux is mediated by alternative or redundant channels. By analogy with *S. cerevisiae*, the *C. neoformans* Pfk2 homolog (CNAG_04076) may also be a Hog1 target. Supporting this metabolic dysregulation model, we observed markedly elevated basal glycerol levels in the membrane-tethered Hog1 strain and its extreme sensitivity to fludioxonil, which hyperactivates the HOG pathway. Future phosphoproteomic analyses will be required to identify the relevant cytoplasmic targets.

The membrane-tethered Hog1 strain was also more sensitive to the ER stressor tunicamycin than the *hog1*Δ mutant, whereas nuclear Hog1 restored resistance. Notably, Ire1-Hxl1-dependent UPR activation (Cheon et al. 2014) was comparable across all strains, indicating that Hog1 and the canonical UPR operate in parallel in *C. neoformans*. This contrasts with *S. cerevisiae*, where membrane-tethered and nuclear Hog1 confer tunicamycin resistance independently of glycerol synthesis (Garcia-Marques et al. 2015), and with *C. albicans*, where Hog1 directly modulates *HAC1* splicing (Husain et al. 2021). Thus, the pronounced ER stress sensitivity of the membrane-tethered Hog1 strain in *C. neoformans* likely reflects a combination of excessive glycerol accumulation and perturbation of an as-yet-uncharacterized non-canonical ER stress response regulator.

Beyond osmotic and ER stresses, we systematically examined additional Hog1-regulated phenotypes (Bahn et al. 2005). For oxidative stress responses, membrane-tethered Hog1 fully restored resistance, while nuclear Hog1 provided only partial restoration, indicating predominant and auxiliary roles for cytoplasmic and nuclear Hog1, respectively—again distinct from *C. albicans*, where either Hog1 pool suffices (Day et al. 2017). Induction of *SRX1*, *TSA3*, *TRX1*, and *TRX2* required both Hog1 pools, implying cooperative action of cytoplasmic and nuclear transcription factors, or direct kinase-mediated regulation of metabolic enzymes. Additionally, both Hog1 pools contributed to thermal and membrane stress resistance, ergosterol biosynthesis, and capsule, melanin, and urease production, though with varying relative contributions. In contrast, cytoplasmic Hog1 played a more prominent role in repressing sexual differentiation: pheromone gene induction and expression of key mating regulators *ZNF2* and *SXI1*α were strongly suppressed by cytoplasmic Hog1 but markedly elevated in the *hog1*Δ mutant. This suggests that a yet-unidentified transcription factor(s) is retained in the cytoplasm through Hog1-mediated phosphorylation, thereby preventing induction of mating-related genes.

The spatially specific Hog1 phenotypes observed here are most consistent with a model in which Hog1 rewires gene expression by coordinately regulating multiple transcription factors. In *S. cerevisiae*, Hog1 interacts with Sko1 (Proft and Struhl 2002), Hot1 (Alepuz et al. 2001; Bai et al. 2015), Smp1 (de Nadal et al. 2003), and Msn2/Msn4 (Capaldi et al. 2008) to drive osmoadaptive transcription, and Sko1 is similarly regulated in *C. albicans* (Su et al. 2013). However, these orthologs are not conserved in *C. neoformans*, suggesting lineage-specific diversification of Hog1 interaction networks. Instead, Atf1 and Mbs1 have been proposed as Hog1-dependent transcription factors in *C. neoformans* (Bahn and Jung 2013; So et al. 2019; Song et al. 2012), analogous to Sty1-Atf1 regulation in *Schizosaccharomyces pombe* (Wilkinson et al. 1996), SakA/MpkC-AtfA signaling in *A. nidulans* (Day and Quinn 2019), and Hog1-Swi4 regulation in *C. albicans* (Sellam et al. 2019). Consistent with this multi-factor model, transcriptomic analyses indicate that Hog1 modulates over 500 stress-responsive genes, including several transcription factors (Goich et al. 2025; Ko et al. 2009).

Taken together, our findings support a model in which nuclear Hog1 in *C. neoformans* orchestrates stress-induced gene expression through transcription factor–dependent programs, while cytoplasmic Hog1 remodels cellular metabolism and signaling networks that indirectly shape nuclear transcriptional outputs (Fig. 9). This explains why *C. neoformans* requires tightly balanced Hog1 localization to coordinate stress adaptation, development, and virulence. More broadly, although Hog1 is conserved as a central stress-response kinase across fungi, its downstream effectors and regulatory strategies have diversified across lineages. Future phosphoproteomic analyses using the Hog1 localization variants generated here will be essential to define the spatial and temporal mechanisms by which Hog1 engages its targets.

## Data availability statement

Supplementary Tables 1 and 2 list the strains and primers used in this study, respectively, and Supplementary Dataset 1 contains the raw fluorescence intensity data underlying the line-scan profiles shown in Fig. 1d. All datasets presented in this study are available in GENETICS online.

## Author Contributions

YB: Conceptualization, Writing – original draft, Writing – review & editing, Methodology, Formal analysis, Validation, Data curation. YS: Conceptualization, Writing – original draft, Writing – review & editing, Investigation, Supervision, Funding acquisition, Project administration.

