## Supplementary material for "Deciphering subcellular localization-dependent functions of Hog1 MAPK in *Cryptococcus neoformans*": Suplemental Figure 1

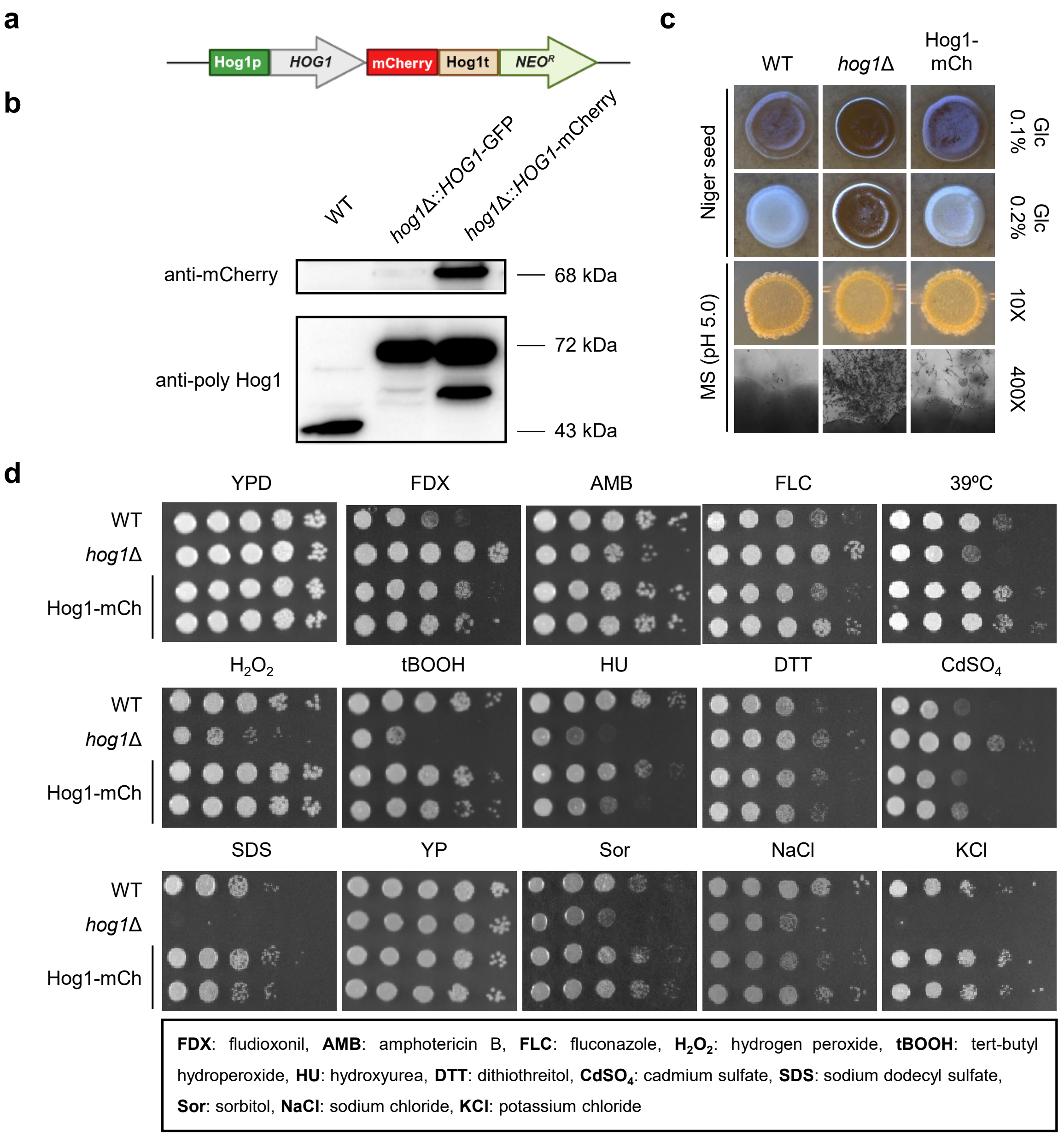


**Supplementary Figure S1. The mCherry tagged chimeric Hog1 mutant restores all altered phenotypes of *hog1*Δ.** (a) The design of the mCherry tagged Hog1 mutant is illustrated by Biorender (https://BioRender.com/v91v944). The native Hog1 promoter and terminator were used to generate the mCherry tagged mutant. (b) The expected protein sizes of Hog1-GFP (YSB242) and Hog1-mCh (YSB7160) were confirmed by western blot analysis using anti-mCherry or polyclonal anti-Hog1 antibodies. (c) Hog1-mCh restores excessive melanin production on niger seed medium and hyperfilamentation on Murashige and Skoog (MS) medium. (d) Two independent Hog1-mCh mutants (YSB7160 and YSB7161) restore various reported *hog1*Δ phenotypes, including sensitivity to antifungal drugs (fludioxonil, amphotericin B, fluconazole), oxidative stresses (hydrogen peroxide, *tert*-butyl hydroperoxide), genotoxic stresses (hydroxyurea, dithiothreitol), heavy metal stress (cadmium sulfate), cell membrane stress (sodium dodecyl sulfate), and osmotic stresses (sorbitol, sodium chloride, potassium chloride).
