## Supplemental Figure 2 for "Deciphering subcellular localization-dependent functions of Hog1 MAPK in *Cryptococcus neoformans*"

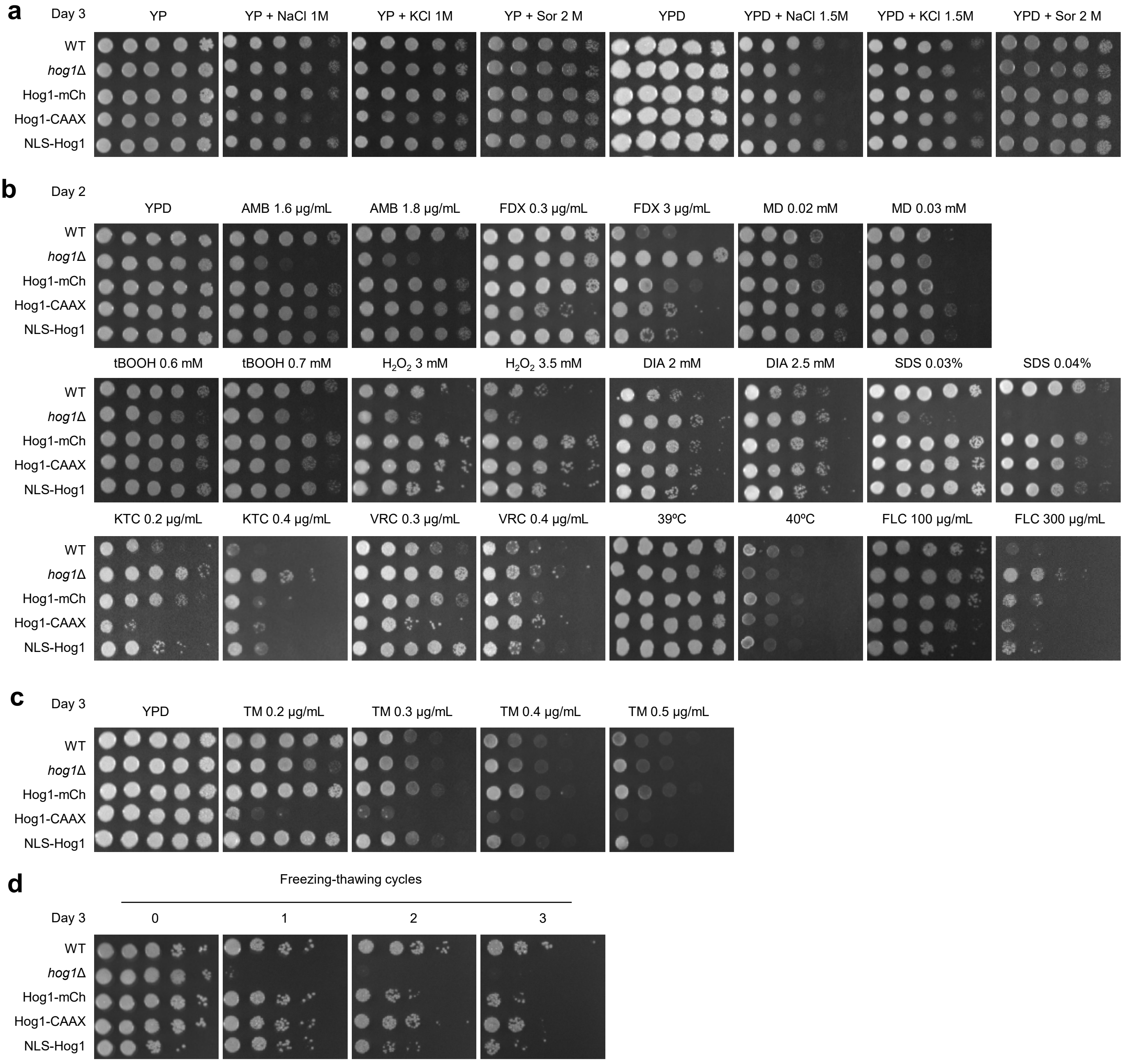


**Supplementary Figure S2. Replicate spotting assays evaluating the phenotypes of *HOG1* mutants under diverse stress conditions.** The phenotypes of wild-type, *hog1*Δ, Hog1-mCh, Hog1-CAAX, and NLS-Hog1 strains were confirmed by duplicate spotting assays. (a) Growth on YP and YPD media supplemented with osmotic stressors (sodium chloride, potassium chloride, and sorbitol [Sor]). (b) Growth on YPD medium under various stress conditions, including antifungal drugs (AMB, amphotericin B; FDX, fludioxonil; FLC, fluconazole; KTC, ketoconazole; VRC, voriconazole), oxidative stressors (tBOOH, tert-butyl hydroperoxide; $H_2O_2$, hydrogen peroxide; DIA, diamide; MD, menadione), a membrane-perturbing agent (SDS, sodium dodecyl sulfate), and heat stress (39°C and 40°C). (c) Growth on YPD medium supplemented with the endoplasmic reticulum (ER) stress inducer tunicamycin (0.2–0.5 μg/mL). Plates were imaged after 3 days of incubation in (a–c). (d) The experiment shown in Fig. 4c was independently repeated using the same strains and conditions (0 to 3 cycles of freezing in liquid nitrogen followed by thawing, then spotting onto YPD medium). Plates were imaged after 3 days of incubation.
