## Supplementary Figure 3 for "Deciphering subcellular localization-dependent functions of Hog1 MAPK in *Cryptococcus neoformans*"

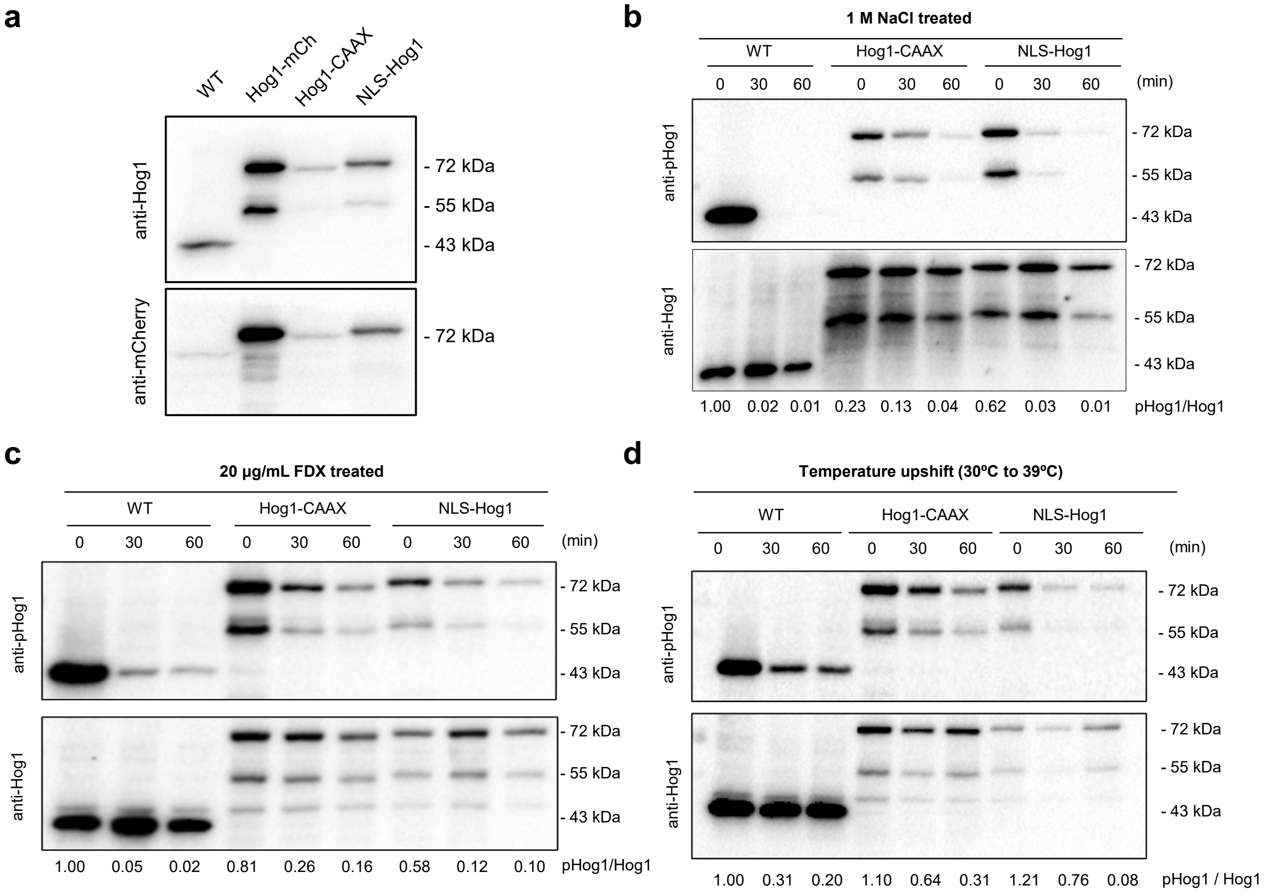


**Supplementary Figure S3. Western blot analysis of Hog1 expression and phosphorylation in wild-type (WT) and Hog1 localization-variant strains.** (a) Basal expression of Hog1 in WT, Hog1-mCh, Hog1-CAAX, and NLS-Hog1 strains under unstressed conditions, detected with anti-Hog1 and anti-mCherry antibodies, corresponding to the biological replicate of Fig. 1b. (b–d) Independent biological replicate experiments corresponding to the western blot data shown in Fig. 2b, Fig. 3b, and Fig. 4b, respectively. The indicated strains (WT, Hog1-CAAX, and NLS-Hog1) were exposed to (b) 1 M NaCl, corresponding to Fig. 2b; (c) 20 µg/mL fludioxonil (FDX), corresponding to Fig. 3b; and (d) heat shock (30°C to 39°C), corresponding to Fig. 4b. Following treatment, total proteins were prepared, and the levels of total and phosphorylated Hog1 were determined by western blotting using anti-Hog1 and anti-phospho-Hog1 (anti-p-p38) antibodies. Signal intensities were quantified with Image Lab software, and the relative phosphorylation levels of Hog1 were calculated by normalizing to the total Hog1 protein levels. In all three stress conditions, Hog1-CAAX consistently sustained higher phospho-Hog1/Hog1 ratios than NLS-Hog1 throughout the time course, in agreement with the main figures and supporting the reproducibility of localization-dependent Hog1 dephosphorylation kinetics.
