## Supplementary Figure 4 for "Deciphering subcellular localization-dependent functions of Hog1 MAPK in *Cryptococcus neoformans*"

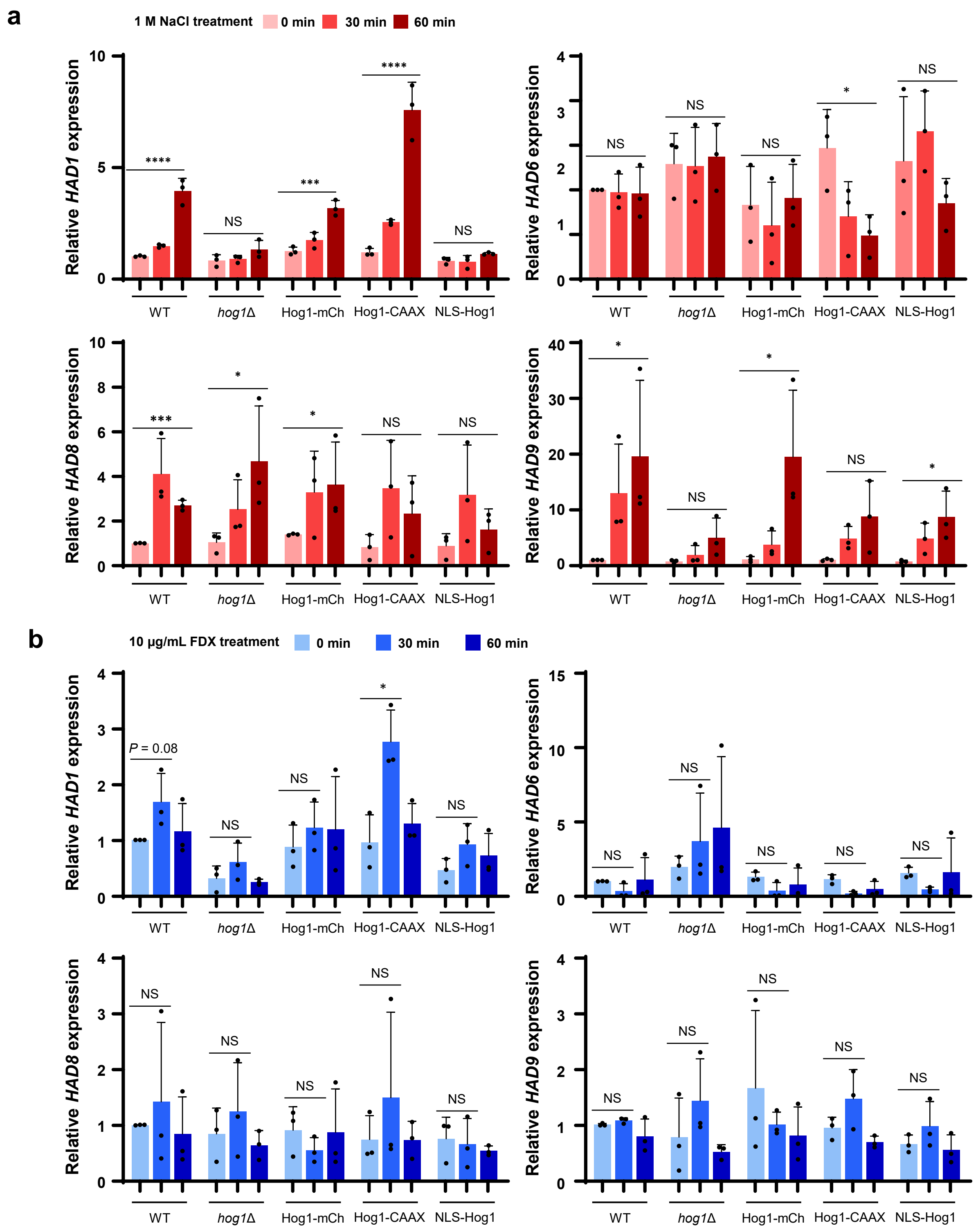


**Supplementary Figure S4. mRNA expression of glycerol-3-phosphatases in *C. neoformans* Hog1 mutants.** (a) Wild type and Hog1 strains (*hog1*Δ, Hog1-mCh, Hog1-CAAX, NLS-Hog1) were treated with 1 M NaCl and sampled at the indicated time points. Transcript levels of *HAD1*, *HAD6*, *HAD8*, and *HAD9* were quantified by qRT-PCR. (b) The same strains were treated with 10 μg/mL fludioxonil and sampled at the indicated time points. *HAD1*, *HAD6*, *HAD8*, and *HAD9* transcript levels were quantified by qRT-PCR. Statistical analysis is performed with two-tailed *t*-test. * indicates $P < 0.05$; ** indicates $P < 0.01$; *** indicates $P < 0.001$; **** indicates $P < 0.0001$; NS indicates a non-significant difference.
