## Supplementary Table 1 for "Deciphering subcellular localization-dependent functions of Hog1 MAPK in *Cryptococcus neoformans*"

**Supplementary Table 1. List of strains used in this study.**

| Strains (YSB #) | Genotype | Parent | Reference |
|---|---|---|---|
| H99 | *MAT*α (Serotype A) | | (Perfect et al. 1993) |
| YL99 | *MAT***a** (Serotype A) | | (Semighini et al. 2011) |
| YSB64 | *MAT*α *hog1*Δ::*NAT*-STM #177 | H99 | (Bahn et al. 2005) |
| YSB7160 | *MAT*α *hog1*Δ::*HOG1-NEO-mCherry* | YSB64 | This study |
| YSB7161 | *MAT*α *hog1*Δ::*HOG1-NEO-mCherry* | YSB64 | This study |
| YSB8841 | *MAT*α *hog1*Δ::*HOG1-mCherry-CaaX-NEO* | YSB64 | This study |
| YSB11710 | *MAT*α *hog1*Δ::$NLS_{SV40}$-*HOG1-mCherry-NEO* | YSB64 | This study |
