## Supplementary Table 2 for "Deciphering subcellular localization-dependent functions of Hog1 MAPK in *Cryptococcus neoformans*"

**Supplementary Table 2. List of primers used in this study.**

| Primer Name | Sequence (5' to 3') | Comments |
|---|---|---|
| **B79** | TGTGGATGCTGGCGGAGGATA | Screening primer |
| **B10048** | AAGCAACAAGGAGTAAGCG | *HOG1*-mCherry complementation screening CSO |
| **B10049** | GATATCAGAGTGAAGACAAAAGGCG | *HOG1*-mCherry complementation CLP |
| **B10083** | TAAAGAACCACTTGGGGG | *HOG1*-mCherry sequencing Seq1 |
| **B10084** | GTTTTTCAACCCAGCGTC | *HOG1*-mCherry sequencing Seq2 |
| **B20733** | GATGCATGCTCGAGCGGCCGCGTGAAG ACAAAAGGCGTG | $NLS_{SV40}$-*HOG1*-mCherry forward primer LP |
| **B20734** | TATCCATCACACTGGCGTACCAATCTAT CCCTCTCTC | $NLS_{SV40}$-*HOG1*-mCherry reverse primer RP |
| **B20891** | GACCTTCCGCTTCTTCTTAGGCATGGTC TATACTGTTAAG | *NLS-HOG1-mCherry* primer L2 |
| **B20892** | GAAGCGGAAGGTCGAGGACCCTATGGC CGATTTTGTCAAG | *NLS-HOG1-mCherry* primer R1 |
| **B11801** | GACGAGCTGTACAAGTGCTGTAGAGGA TGCGTCGTCCTCTAACTCGAG | *HOG1-CAAX-mCherry* primer R1 |
| **B11643** | GCACCCGAGATCATGTTGAC | *HOG1-CAAX-mCherry* primer seq3 |
| **B10727** | TGAACTCCTTGATGATGGC | mCherry sequencing primer |
| **B2861** | GATGAAGCACTTTTGCCAAG | *GPD1* qRT primer qLP |
| **B2862** | AGAAAGACACGAAGTAATCA | *GPD1* qRT primer qRP |
| **B22079** | GGCTATGGGTCAATTCTGCG | *GPD2* qRT primer qLP |
| **B22080** | CATTGACCCGGAAAGAAGGC | *GPD2* qRT primer qRP |
| **B22081** | TCCATTTCCTCGGTCTTGCT | *ENA1* qRT primer qLP |
| **B22082** | TGTCAACATTCGAGGGGTGA | *ENA1* qRT primer qRP |
| **B22085** | GCCATTCCCTGACCCTTACT | *HAD1* qRT primer qLP |
| **B22086** | CGCCTTGCCTGATCGAATAC | *HAD1* qRT primer qRP |
| **B22847** | TGCTGGCAAGGAAGGTCTTA | *HAD6* qRT primer qLP |
| **B22848** | TAGACAGTGGTTGCAGACGT | *HAD6* qRT primer qRP |
| **B22089** | TTGCACCAGTCACAAAAGGG | *HAD8* qRT primer qLP |
| **B22090** | AACTTCGTCCCCTTCAGTGT | *HAD8* qRT primer qRP |
| **B22091** | TCATGGATGCCTGAACCGTA | *HAD9* qRT primer qLP |
| **B22092** | AAGAAGGGGTCAGGGAAAGG | *HAD9* qRT primer qRP |
| **B6715** | GGTTCAACGTCAGTAATCCAAGTG | *SRX1* qRT primer qLP |
| **B1697** | GGCAGGAACATCAATAATCC | *SRX1* qRT primer qRP |
| **B10132** | CGACTACTGGGCGACTTG | *TRX1* qRT primer qLP |
| **B10133** | GCAAACTTGACATTGGGG | *TRX1* qRT primer qRP |
| **B10134** | CACGCTTCTCCTATTTCG | *TRX2* qRT primer qLP |
| **B10135** | GTCTACCAAAACGGGTCTG | *TRX2* qRT primer qRP |

| | | |
|---|---|---|
| **B10136** | GATGAGGATAGGACAGTG | *TSA3* qRT primer qLP |
| **B10137** | GAAAACAGTCCTGACCGTAAAC | *TSA3* qRT primer qRP |
| **B10138** | GAGGCACCTTCTTCATTG | *TSA1* qRT primer qLP |
| **B10139** | TTGATGACACGGATGGTC | *TSA1* qRT primer qRP |
| **B5251** | CACTCCATTCCTTTCTGC | *HXL1* splicing forward primer |
| **B5252** | CGTAACTCCACTGTGTCC | *HXL1* splicing reverse primer |
| **B9062** | GTTCGAGACTTTCAATGCCC | *ACT1* qRT primer qLP |
| **B9063** | ACCAGAGTCAAGAACGATAC | *ACT1* qRT primer qRP |
| **B8097** | CGCCTTCACTGCCATCTTC | *MFα1* qRT primer qLP |
| **B8098** | ACAAAGGGTCATGCCACCGG | *MFα1* qRT primer qRP |
| **B8557** | CTGCGATTTCGGTCTTGCC | *CPK1* qRT primer qLP |
| **B8558** | GATACCACCTTGTAGCGAC | *CPK1* qRT primer qRP |
| **B8555** | CCATCAGGTAGCAAAGTAGC | *CPK2* qRT primer qLP |
| **B8556** | CAAAGTACTTCAACAGCTT | *CPK2* qRT primer qRP |
| **B11381** | TGAGGGAGAAAAGGTTCC | *MAT2* qRT primer qLP |
| **B11382** | GGAGGAGGCATTGACTTATTC | *MAT2* qRT primer qRP |
| **B11060** | TTGTGAACGACCACAACCAC | *ZNF2* qRT primer qLP |
| **B11061** | TGCCTTGCAAGATCACTTTTT | *ZNF2* qRT primer qRP |
| **B19004** | GCAATTCACCAAAGCCCTCA | *SXI1α* qRT primer qLP |
| **B19005** | CTTTCTGTTGTGCGGTTGGA | *SXI1α* qRT primer qRP |
